# Functional Type I and Type II interferon crosstalk restricts progenitor exhausted CD8 T cells through spatial exclusion and checkpoint enforcement

**DOI:** 10.64898/2026.03.01.708869

**Authors:** Siran Liu, Heidi J. Elsaesser, Rene Quevedo, Diala Abd-Rabbo, Bruna C. Bertol, Wenxi Xu, Melissa Yi Ran Liu, Sabelo Lukhele, Sara Lamorte, Tracy L. McGaha, David G. Brooks

## Abstract

Type I interferon (IFN-I) and interferon-γ (IFNγ) are central regulators of antiviral immunity, yet how they cooperatively govern CD8 T cell fate during chronic infection remains unresolved. Here, we uncover a previously unrecognized, spatially encoded interferon circuit that actively constrains progenitor exhausted CD8 T cells (Tpex) during chronic LCMV infection. Persistent IFN-I signaling indirectly restricts Tpex expansion by enforcing their sequestration within PDL1–rich B cell niches of lymphoid tissue and by suppressing T cell-derived IFNγ. Blockade of IFN-I signaling enables Tpex migration into T cell zones of splenic follicles driving IFNγ production, which in turn sustains PDL1 expression on myeloid cells to re-impose local inhibitory pressure. Combined IFN-I and IFNγ blockade disrupts this feedback, promoting coordinated niche redistribution of Tpex and checkpoint remodeling that drives robust Tpex expansion. Single-cell transcriptomics reveal that this layered IFN-I–IFNγ interplay establishes a regulatory balance that constrains Tpex proliferation while preserving effector-like transcriptional programs in their progeny effector CD8 T cells, ultimately preventing premature terminal differentiation. Thus, interferons orchestrate the coordinated T cell–myeloid regulatory circuit that integrates tissue organization, cytokine feedback, and checkpoint control to regulate CD8 T cell exhaustion during chronic infection.

## INTRODUCTION

Type I interferon (IFN-I) and type II interferon (IFNγ) are well-established inhibitors of viral replication and mediators of immune activation^**1,2**^. However, their functions are multifaceted, coordinating immune stimulation with suppressive mechanisms that limit immunopathology^**2**^. During chronic viral infection, IFN-I signaling exerts strong inhibitory effects that prevent viral clearance^**3,4**^. Paradoxically, blockade of IFN-I signaling at the onset of chronic infection enhances immune cell function and accelerates viral clearance, revealing a negative regulatory role for IFN-I signaling^**2,3**^. IFN-I blockade also increases IFNγ expression, and this rise in IFNγ is required for the improved viral control, indicating the presence of a tightly regulated feedback circuit between IFN-I and IFNγ signaling^**3**^.

During chronic viral infection, persistent antigen exposure together with a systemically immunosuppressive environment drives progressive attenuation of CD8 T cell function, a state termed exhaustion, ultimately limiting viral control^**5**^. T cell exhaustion is an intrinsic differentiation program maintained through coordinated transcriptional, epigenetic, and metabolic reprogramming^**5–8**^. This state is characterized by diminished effector and cytolytic activity, sustained expression of inhibitory receptors (IRs; PD1, CD39, Lag3, Tim3), and reduced proliferative capacity^**5–8**^. Recently, a subset of precursor exhausted CD8 T cells (Tpex) with self-renewing potential has been identified in both chronic viral infection and cancer^**9**^. Tpex cells proliferate in response to incompletely defined cues to sustain downstream exhausted effector (Teff) and terminally exhausted (Tex) T cell populations during chronic infection^**9–11**^. The Tpex population is therapeutically relevant, as Tpex cells preferentially respond to PD-1/PD-L1 blockade in both chronic infection and cancer^**9,12**^. Importantly, IFN-I signaling inhibits Tpex differentiation at the onset of chronic infection, suggesting that IFN-I constrains the establishment of the Tpex pool and biases the response toward terminal Tex fates^**13**^.

At the onset of what will become a chronic infection, CD8 T cells are primed within a highly inflammatory environment that progressively transitions into an immunosuppressive state, ultimately promoting T cell exhaustion^**3**^. In an established chronic infection, however, this suppressive milieu is already entrenched. By this stage, T cell differentiation and the transcriptional and epigenetic programs underpinning exhaustion are largely fixed, and T cells function within a profoundly altered landscape characterized by suppressive dendritic cells and macrophages, persistent antigen exposure, and sustained TCR signaling^**14**^. The role of interferons in shaping T cell differentiation and effector function during early priming has been extensively studied^**15,16**^. However, how interferons continue to regulate T cell responses once chronic infection is established remains poorly defined. This distinction is clinically consequential, as interventions delivered at disease onset aim to prevent exhaustion, whereas therapies administered during established infection or cancer must instead reinvigorate already exhausted T cells, a considerably greater challenge. Notably, blockade of IFN-I signaling during established chronic infection lowers viral titers^**3,4**^, indicating that IFN-I continues to impose immune restraint during ongoing disease, yet the mechanisms underlying this suppression have remained unresolved.

Here, we uncover a spatially organized and cytokine-coupled interferon circuit that sustains CD8 T cell suppression during established chronic LCMV infection. In contrast to its role during early T cell priming^**13,16**^, persistent IFN-I signaling indirectly limits the proliferation of Tpex and exhausted effector CD8 T cells once chronic infection is established. This suppression occurs through two coordinated mechanisms. First, IFN-I restrains T cell–derived IFNγ, such that blockade of IFN-I unleashes IFNγ production that paradoxically continues to limit Tpex expansion. Second, IFN-I enforces the spatial confinement of Tpex within PD-L1–rich B cell regions of lymphoid tissue, restricting their interactions with dendritic cells and macrophages and thereby maintaining suppression. When IFN-I signaling is relieved, Tpex migrates into T cell zones and initiates expansion. However, the concurrent rise in IFNγ acts on myeloid cells to sustain PD-L1 expression and re-establish local inhibitory pressure. As a result, robust Tpex proliferation and downstream effector generation occur only when both IFN-I and IFNγ signaling are blocked, enabling coordinated spatial redistribution together with checkpoint remodeling. Collectively, these results define the interferon network as a multilayered suppressive system that integrates tissue organization, cytokine feedback, and checkpoint control to constrain Tpex expansion and stabilize exhaustion.

## RESULTS

### IFN-I and IFNγ signaling during the established chronic infection suppress virus-specific CD8 T cell proliferation

To investigate how prolonged IFN-I and IFNγ signaling shape CD8 T cell differentiation and exhaustion during chronic LCMV infection, we adoptively transferred LCMV-glycoprotein (GP)_33-41_-specific TCR transgenic CD8 T cells (P14 cells) into naïve mice, followed by infection with LCMV-Cl13 to establish chronic infection. Beginning at day 25 post infection (once the chronic infection is firmly established), mice received antibody treatments to block IFN-I signaling (anti-IFNAR), IFNγ signaling (anti-IFNγ), or combined IFNAR and IFNγ double blockade (DB) (Fig. S1A). CD8 P14 T cells were analyzed one day after the final antibody administration (35 days after infection) using time-of-flight mass cytometry (CyTOF). Consistent with prior observations, plasma viral titers increased immediately after the final anti-IFNAR treatment (Fig S1B). In contrast, IFNγ blockade alone did not alter viral titers, whereas combined IFN-I and IFNγ double blockade (DB) resulted in higher viral loads than IFNAR blockade alone (Fig S1B). These findings indicate that IFN-I signaling continues to restrain viral replication during chronic infection and that, upon IFN-I blockade, IFNγ acquires a previously masked antiviral role.

CyTOF analysis identified 11 clusters within the CD8 P14 compartment that segregated into three major differentiation states^**7,14**^. Tpex cells localized to clusters (c)1 and c2 (TCF1+, CD39-, CD69+/-); exhausted effector (Teff) cells to c3, c5, c6 and c7 (TCF1-, CX3CR1+, CD39+, CD69-); and terminally exhausted (Tterm) cells to clusters c8–c11 (TCF1⁻ CX3CR1⁻ CD39⁺ CD69⁺) (Fig. 1A, B). Within the Teff compartment, clusters c3 and c7 were proliferating (Ki67⁺) and corresponded to intermediate exhausted (Tint) cells representing the immediate progeny of Tpex.

**Figure 1.**
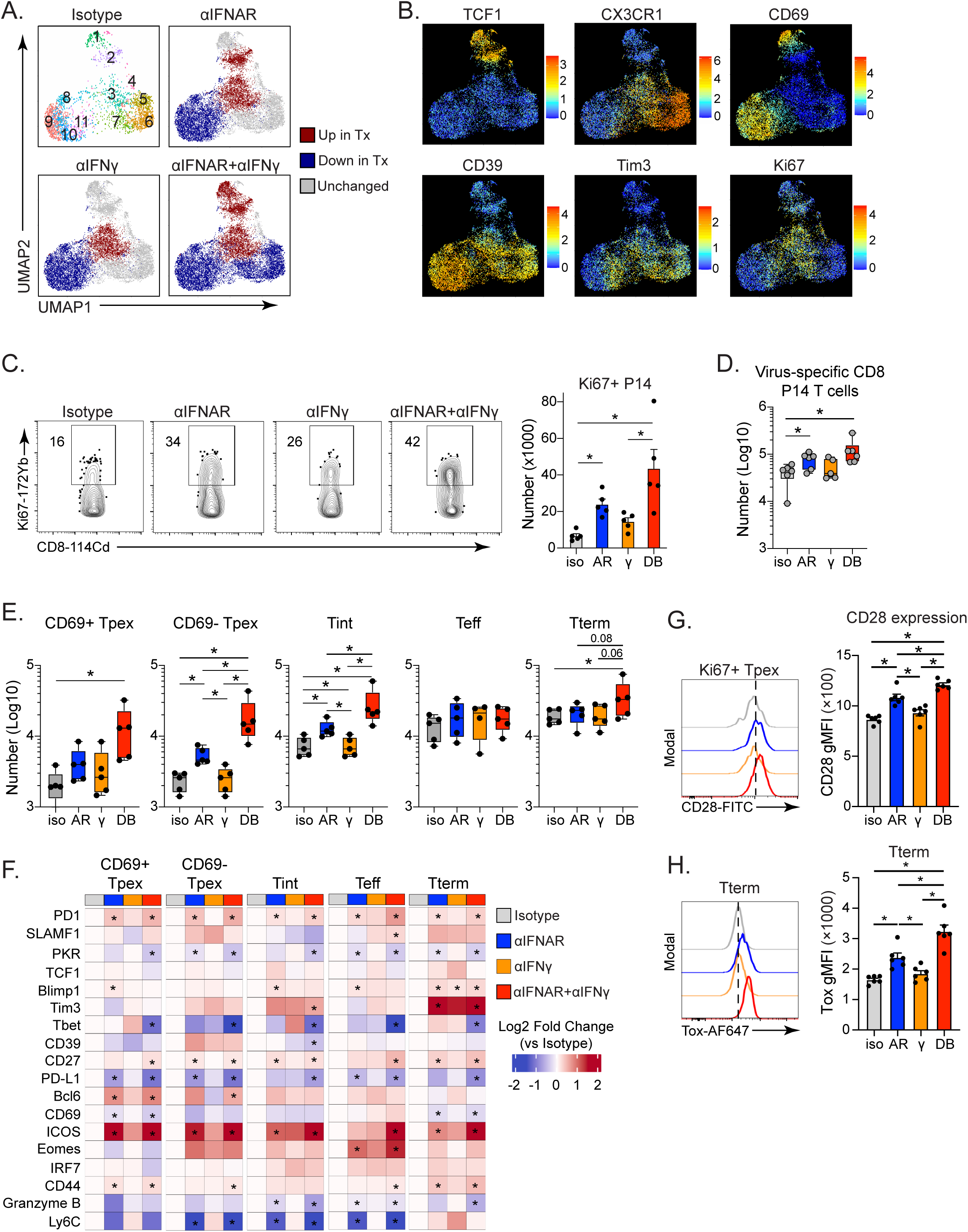
Increased Tpex following anti-IFNAR and anti-IFNAR+anti-IFNγ double blockade. Mice were treated with isotype (iso), anti-IFNAR (AR), anti-IFNγ (γ) or anti-IFNAR+IFNγ double blocking (DB) antibodies beginning on day 25 after LCMV-Cl13 infection. Data in the figure are from the spleen one day following the final antibody treatment (day 35 after infection). **(A)** UMAPs show virus-specific CD8 P14 T cell Phenograph clusters of CyTOF after the indicated antibody treatment. Clusters with significantly increased abundance (red) or decreased abundance (blue) following treatment (Tx). Clusters in gray have no significant proportion change. **(B)** UMAPs show the expression of the indicated subset-defining protein in the virus-specific CD8 P14 T cell clusters. **(C)** Flow plots show the percent and the bar graph shows the number of Ki67-positive virus-specific CD8 P14 T cells following each treatment. **(D, E)** Box plots show the number of **(D)** virus-specific CD8 P14 T cells and **(E)** of the indicated Tpex and Tex populations. **(F)** The heatmaps show the Log2 fold-change in protein expression (left) in the indicated virus-specific CD8 P14 T cell subset (above each heatmap) following each blockade. **(G, H)** Histograms show the expression and bar graphs the geometric mean fluorescence intensity (gMFI) of **(G)** CD28-expressing Tpex, and **(H)** Tox-expressing Tterm populations of virus-specific CD8 P14 T cells. Data are representative of 3-5 experiments with 5-6 mice per group. Circles indicate individual mice. The line in each box of the box plots indicates the median, and the bars indicate the highest and lowest value. *, p<0.05 by one-way ANOVA.

At day 35 after LCMV-Cl13 infection, the frequency and number of proliferating (Ki67⁺) CD8 P14 T cells increased following IFNAR blockade and double blockade, whereas IFNγ blockade alone had no effect (Fig 1C). Consistent with this enhanced proliferation, total CD8 P14 cell numbers increased under IFNAR blockade and double blockade (Fig. 1D). IFNAR single blockade significantly increased the frequency and number of the CD69- Tpex subset (i.e., the cycling pool of Tpex, Fig 1A, B), with the most pronounced effect observed following combined IFNAR and IFNγ blockade, whereas blocking IFNγ signaling alone did not affect the frequency or number of CD69- Tpex (Fig.1E, Fig. S1C). Expansion of the proliferatively quiescent CD69+ Tpex population^**7**^, was only observed with dual IFN-I and IFNγ blockade (Fig. 1E, Fig. S1C). The increase in Tpex was accompanied by expansion of the downstream Tint population following IFNAR blockade and double blockade (Fig. 1E). Tint cells also showed a moderate increase after IFNγ blockade alone; however, this did not translate into increased total CD8 P14 cells (Fig. 1D, E). The number of Teff and Tterm populations remained stable across all treatments (Fig. 1E), suggesting that newly expanded Tpex did not preferentially progress toward terminal exhaustion. Phenotypic analysis revealed broad upregulation of activation and exhaustion-associated molecules, including PD1, ICOS, CD44, and CD27, across Tpex and Tex populations following IFNAR blockade and double blockade (Fig. 1F), consistent with a heightened activation state. Minimal phenotypic changes were detected following IFNγ blockade alone (Fig. 1F). Within proliferative Tpex, CD28 expression was selectively increased after IFNAR blockade and double blockade (Fig. 1G), suggesting enhanced responsiveness to co-stimulatory signals relevant for anti-PD1 therapy^**17**^. Although Tex populations were not numerically expanded by interferon blockade, their phenotype was altered. Effector-associated molecules such as T-bet and Granzyme B were reduced following IFNAR blockade and double blockade (Fig. 1F). Additionally, Tim3+ Tterm cells increased across all three treatments, with the strongest effect under double blockade, while Tim3⁻ Tterm and total Tterm numbers remained unchanged (Fig. 1F, Fig. S1D), indicating de novo Tim3 upregulation and progression toward deeper exhaustion^**10**^. Consistent with this, the exhaustion-associated transcription factor Tox was significantly upregulated within Tterm following IFNAR blockade and double blockade (Fig. 1H). Finally, although cytokine production in Tpex remained unchanged, IFNAR blockade and double blockade reduced the frequency of polyfunctional IFNγ⁺ TNFα⁺ Tex cells (Fig. S1E). Thus, IFN-I signaling restrains Tpex expansion while simultaneously supporting effector features within Tex populations during established chronic infection.

### IFN-I and IFNγ signaling indirectly suppress Tpex expansion

We next asked whether interferon signaling directly suppresses Tpex formation during established chronic infection. Analysis of receptor expression revealed that IFNγR was readily detected on Tex but was absent on Tpex and remained undetectable following IFNAR blockade (Fig. 2A), providing a potential explanation for why IFNγ blockade alone did not enhance Tpex proliferation or expansion.

**Figure 2.**
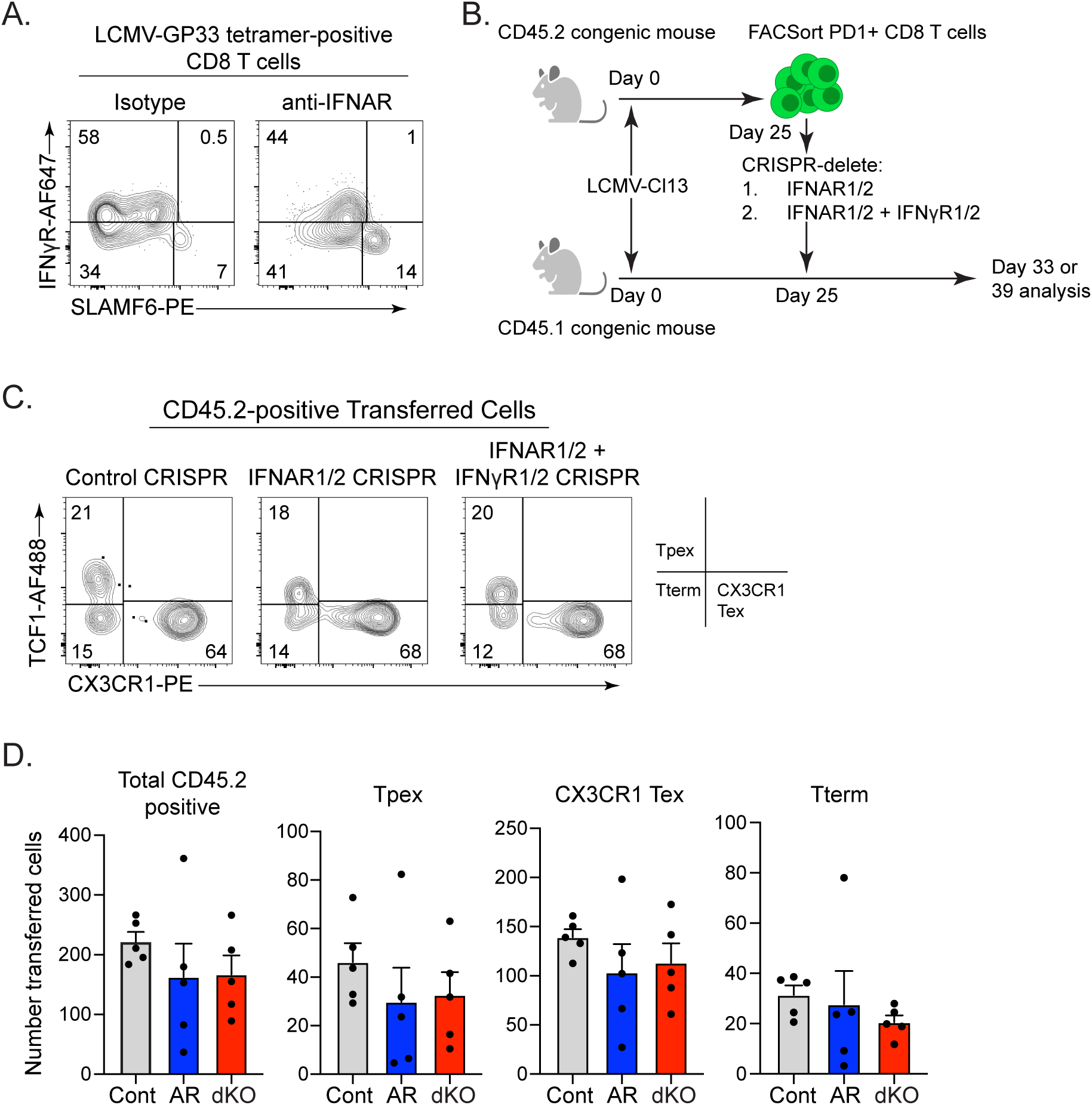
IFN-I and IFNγ indirectly control Tpex proliferation. **(A)** Flow plots show IFNγR1 expression on LCMV-GP33-41 tetramer-positive CD8 T cells in the spleen following isotype or anti-IFNAR blockade (day 33 after LCMV-Cl13 infection). Tpex are SLAMF6-positive. **(B)** Experimental schematic for the CRISPR experiments: CD45.1 (recipient) and CD45.2 (donor) mice were infected with LCMV-Cl13. Twenty-five days later, PD1+ CD8 T cells were FACSorted and treated with gRNAs targeting IFNAR1, IFNAR2, IFNγR1 and IFNγR2. Cells were then transferred into the infection matched CD45.1-positive mice. Analysis was performed 8-14 days later. **(C)** Flow plots show TCF1 vs CX3CR1 expression by the adoptively transferred, CRISPR-deleted PD1+ CD8 T cells. The right plot shows the position of the indicated subset on the flow plots. **(D)** Bar graphs show the number of total CD45.2-positive transferred cells and the indicated CD8 T cell subset eight days after transfer. Cont is control CRISPR, AR = IFNAR1 and IFNAR2 CRISPR deleted, dKO = double knockout, IFNAR1, IFNAR2, IFNγR1, and IFNγR2 CRISPR deleted. Data are representative of 4 experiments with 3-5 mice per group. Circles in the bar graph indicate individual mice. *, p<0.05 by one-way ANOVA.

To directly test cell-intrinsic interferon signaling, we performed ex vivo CRISPR/Cas9-mediated deletion of IFNAR1/2 (targeting both chains of the IFNAR) or combined deletion of IFNAR1/2 and IFNγR1/2 (also targeting both chains of the IFNγR) in exhausted CD8 T cells. Congenic CD45.1 and CD45.2 mice were infected with LCMV-Cl13, and on day 25 post infection, PD1+ CD8 T cells were FACS-purified from CD45.2 donors (Fig. 2B). Immediately following isolation, CRISPR-mediated receptor deletion was performed using guide (g)RNAs targeting IFNAR1/2 or IFNAR1/2 plus IFNγR1/2, with a non-targeting gRNA serving as control (Fig. S2A-C). Edited cells were transferred into infection-matched CD45.1 recipients and analyzed 8 or 14 days later. Efficient IFNAR1 deletion was detected within 24 hours in culture (Fig. S2C). IFNγR1 expression was undetectable after short-term culture but was efficiently deleted in vivo, as confirmed 8 days after transfer (Fig. S2D, S2E). Further, CRISPR-mediated deletion of IFNAR and IFNγR reduced expression of the interferon-stimulated gene (ISG) PKR within transferred CD8 T cells (Fig. S2F), corroborating the CRISPR-mediated loss of intrinsic interferon signaling.

Unexpectedly, deletion of IFNAR alone or in combination with IFNγR did not increase Tpex or Tex populations (Fig. 2C, D). In contrast, transfer of CRISPR-edited cells lacking both IFNAR and IFNγR into infection matched recipients that received systemic anti-IFNAR and anti- IFNγ double blockade increased the total number of transferred cells and expanded the Tpex compartment (Fig S2G). Together, these data indicate that interferon-mediated restraint of Tpex expansion is not CD8 T cell intrinsic. Importantly, no differences in Tpex frequency were observed between CRISPR-edited IFNAR/IFNγR-deficient cells and CRISPR control cells following blockade (Fig. S2G), confirming that interferon-dependent regulation of Tpex expansion operates predominantly through extrinsic mechanisms. Although interferon receptor deletion did not intrinsically alter Tpex or Tex abundance, Thus, while IFN-I and IFNγ directly induce ISG expression within exhausted CD8 T cells, their suppression on Tpex expansion during chronic infection is mediated through CD8 T cell extrinsic mechanisms.

### Single-cell transcriptomics reveals interferon-dependent control of CD8 T cell fate and function

Although interferon signaling did not directly control the numerical expansion of Tpex or Tex populations, exhausted CD8 T cells remained responsive to ongoing interferon cues during chronic infection. To define how persistent interferon signaling shapes CD8 T cell states, we performed single-cell RNA sequencing (scRNA-seq) on CD8 P14 T cells following interferon blockade during LCMV-Cl13 infection. Consistent with CyTOF analysis, scRNA-seq resolved five major clusters with similar shifts in abundance across treatment conditions (Fig. 3A, B; Fig. S3A). Due to limited detection of CD69 transcripts, Tpex could not be resolved by CD69 expression, and cycling cells were grouped together irrespective of lineage identity^**18**^. Two CX3CR1⁺ Tex subsets were identified, CX3CR1⁺ Tint and CX3CR1⁺ Teff, corresponding to populations defined by CyTOF. Accordingly, Tpex were analyzed independent of CD69, while CX3CR1⁺ Tex subsets were examined separately^**14, 19**^.

**Figure 3.**
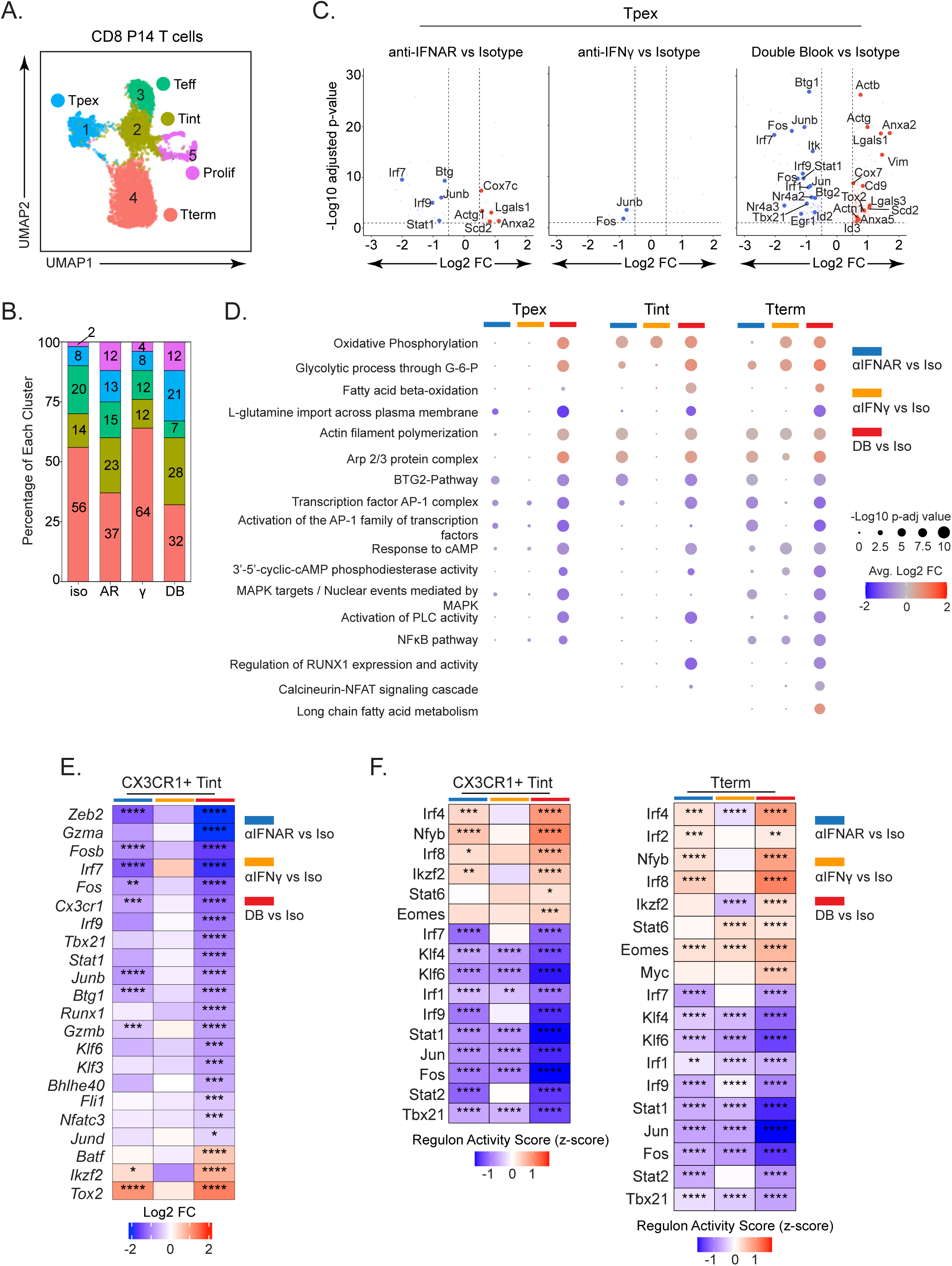
scRNA-seq reveals transcriptional changes in anti-IFNAR and anti-IFNAR+anti-IFNγ double blockade. **(A, B) (A)** UMAPs show virus-specific CD8 P14 T cell clusters in the scRNA-seq and the **(B)** bar graph shows the frequency of each cluster in each of the 4 antibody treatments. **(C)** Volcano plots show the differentially expressed genes in Tpex cells in anti-IFNAR (AR), anti-IFNγ (γ), or combined anti-IFNAR and anti-IFNγ double block (DB) versus isotype control. Red circles equal up and blue circles down compared to isotype control in the indicated antibody treatment group. Cutoff values are set at p_adj < 0.05 and (Log2 FC) > 0.5. **(D)** Pathway analysis indicating the Log2 average fold change (FC) in the Tpex, Tint and Tterm cells following anti-IFNAR, anti-IFNγ, or combined anti-IFNAR and anti-IFNγ double blockade compared to isotype control. **(E)** The heatmap shows differentially expressed genes in CX3CR1+ Tint in anti-IFNAR, anti-IFNγ, or double block versus Isotype control. **(F)** The heatmaps indicate regulon changes (SCENIC analysis) in CX3CR1+ Tint and Terminal Tex in anti-IFNAR, anti-IFNγ, or double block versus Isotype control. * p_adj<0.05, ** p_adj <0.01, *** p_adj <0.001, **** p_adj<0.0001

Differential expression analysis comparing each antibody treatment to isotype controls revealed that Tpex, CX3CR1⁺ Tint, and Tterm populations accounted for most transcriptional remodeling, whereas CX3CR1⁺ Teff cells exhibited minimal changes (Fig. S3B), consistent with their stable abundance across treatments. Double blockade induced the greatest number of differentially expressed genes (DEGs) (Fig. S3B). IFNγ blockade alone produced limited transcriptional effects overall, although 401 DEGs were detected within the Tterm population, suggesting a contribution of IFNγ signaling to transcriptional homeostasis when IFN-I signaling remained intact (Fig. S3B). Given their pronounced phenotypic and numerical changes, subsequent analyses focused on Tpex, CX3CR1+ Tint, and Tterm populations. Across these subsets, downregulation of interferon-stimulated genes (ISGs) and IFN-I signature transcripts was a consistent feature of IFN-I and IFN double blockade, confirming ongoing IFN-I signaling within exhausted CD8 T cells (Fig. 3C; Fig. S3C, S3D). Of note, Tpex displayed the most pronounced reduction in ISGs following double blockade (Fig. 3C; Fig. S3D). Because Tpex lack IFNγ receptor expression (Fig. 2A), the transcriptional changes following double blockade likely reflect a combination of intrinsic IFN-I signaling and altered extrinsic signaling cues within the tissue microenvironment due to IFNγ blockade (Fig. S2F; Fig. S3D).

Among all subsets, Tpex exhibited the strongest IFN-I signaling signature, supporting a direct role for IFN-I in shaping Tpex transcriptional identity (Fig. S3C). Concurrent with ISG downregulation, IFNAR blockade alone and double blockade reduced expression of transcription factors linked to Tpex maintenance and differentiation, including AP-1 family members (Jun, Junb, Fos, Fosb) and additional regulators (Tbx21, Egr1, Stat1, Nr4a2, Nr4a3), suggesting redirection of TCR signaling pathways (Fig. 3C)^**20–22**^. In contrast, genes and pathways associated with oxidative metabolism, cytoskeletal remodeling, and cell motility—including Scd2, Cox6c, Cox7c; Actb, Actg1, Vim; and Cd9, Anxa2, Anxa5, Lgals1, and Lgals3—were upregulated following IFN-I or IFN double blockade (Fig. 3C, D). The enrichment of oxidative phosphorylation and fatty acid metabolism signatures is consistent with increased exhaustion-associated metabolic remodeling following interferon withdrawal^**23,24**^.

To understand how loss of interferon signaling promoted terminal exhaustion rather than effector differentiation, we examined CX3CR1+ Tint cells, the immediate progeny of Tpex. IFNAR blockade, further reinforced by double blockade, markedly reduced expression of transcription factors sustaining effector function in chronic infection, including Zeb2, Tbx21, Fos, Stat1, Btg1, Runx1, Klf6, Klf3, and Bhlhe40 (Fig 3E)^**14,20,25–28**^. These changes suggest that IFN-I, and to a lesser extent IFNγ, normally maintain effector-like transcriptional programs within Tint cells, and that loss of interferon signaling biases differentiation toward terminal exhaustion. To further define regulatory network changes, we applied SCENIC analysis, which identifies transcription factor regulons based on co-expression and cis-regulatory motif enrichment^**29**^. This analysis revealed increased activity of exhaustion-associated regulons (Irf2, Irf4, Irf8, Eomes) alongside reduced activity of effector-supporting regulons (Stat1, Stat2, Klf4, Klf6, Jun, Fos, Tbx21) in CX3CR1⁺ Tint and Tterm populations following IFN-I or IFN double blockade (Fig 3F)^**19,25,30,31**^. Thus, ongoing IFN-I and IFNγ signaling preserve effector-like transcriptional programs and metabolic balance within exhausted CD8 T cells, thereby restraining excessive differentiation of Tint cells into terminally exhausted states.

### Chronic IFN-I signaling restricts Tpex localization in the B cell niche of splenic follicles

Having defined transcriptional programs associated with Tpex expansion, we next examined how interferon signaling indirectly regulates this process through spatial organization. During chronic LCMV infection, Tpex preferentially reside within B cell follicles, a niche characterized by limited antigen availability and reduced dendritic cell and macrophage density that supports their quiescent maintenance^**11,32–34**^. Consistent with this, the Tpex in isotype antibody-treated mice were also enriched in the B cell regions (Fig 4A, B). Strikingly, IFN-I blockade alone, as well as combined IFN-I and IFNγ blockade, significantly increased the proportion of Tpex detected within the T cell zones, with up to ∼40% of Tpex relocating to these regions, with a concordant decrease in the frequency of Tpex in the B cell region of the follicle (Fig 4A, B). In contrast, IFNγ blockade alone did not alter Tpex positioning, as cells remained largely confined to B cell follicles like isotype treatment (Fig. 4A, B). Importantly, the frequency of Tpex present within T cell zones was comparable between IFNAR single blockade and combined IFNAR plus IFNγ blockade (Fig. 4B), indicating that IFN-I signaling is the dominant regulator of Tpex sequestration within the B cell niche and that relief of IFN-I signaling permits their redistribution into T cell zones.

**Figure 4.**
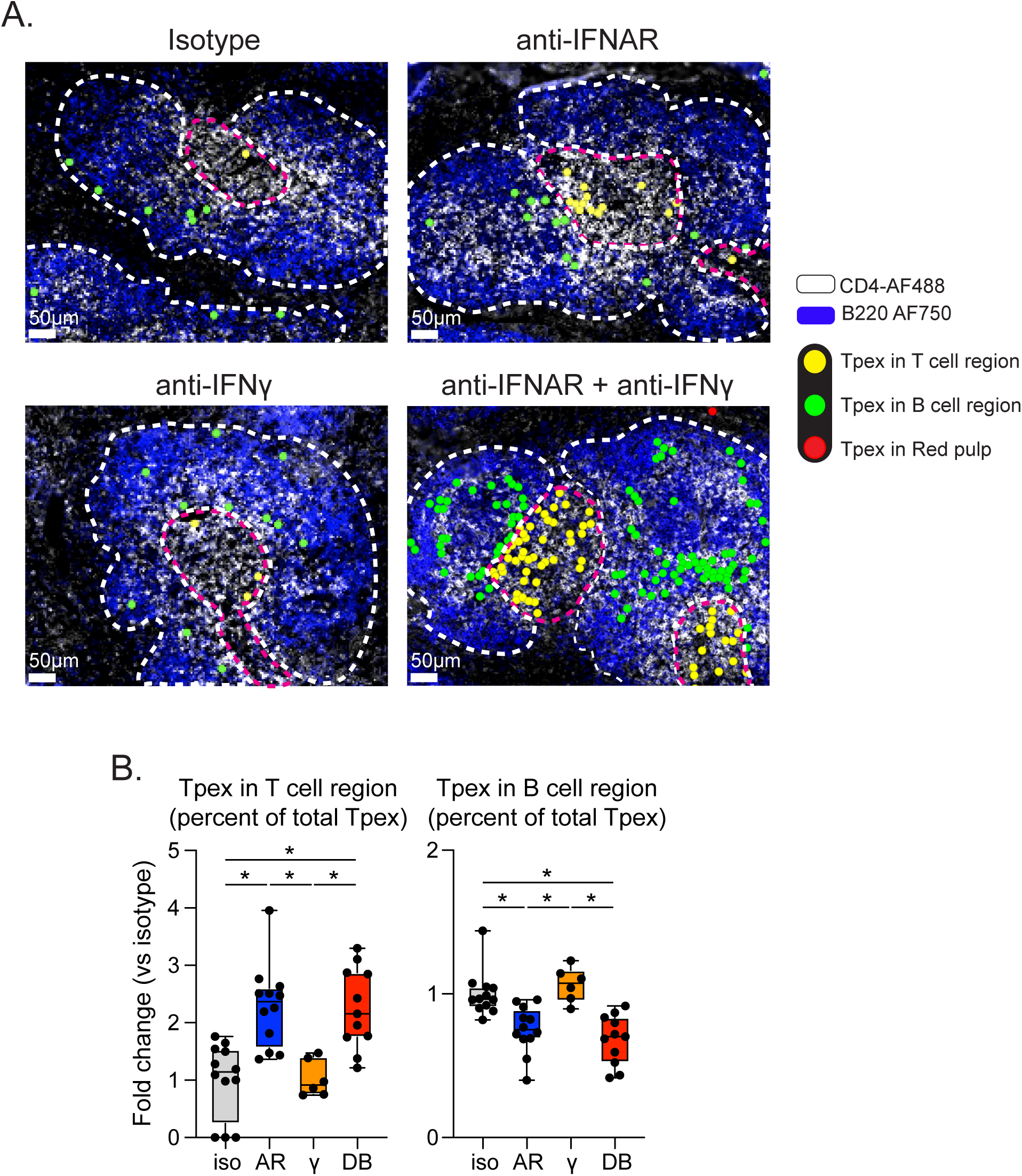
Spatial sequestration of Tpex in the B cell regions by IFN-I. **(A)** Representative immunofluorescence images show Tpex localization in the B cell region (outlined in white) and T cell region (outlined in pink) of the spleen one day after the final antibody treatment. Solid circles indicate: Tpex in the T cell region (yellow), Tpex in the B cell region (green), Tpex in the red pulp (red). **(B)** Bar graphs show the fold change in the percent of Tpex in the follicular T cell regions and B cell regions from whole spleen images in isotype (iso), anti-IFNAR (AR), anti-IFNγ (γ), or combined anti-IFNAR and anti-IFNγ double block (DB). Data is pooled from two individual experiments, with 6-12 spleens per condition. Circles in the bar graph indicate individual mice. * p<0.05, by one-way ANOVA

### IFN-I blockade increases IFNγ expression by T cells to sustain PDL1 expression on myeloid cells

We next examined how IFNγ is modulated following disruption of chronic IFN-I signaling and how this response constrains Tpex and Tex expansion. The minimal effect of blocking IFNγ without IFNAR blockade suggests the presence of an IFN-I-IFNγ feedback circuit. To identify cellular sources of IFNγ, we used IFNγ-YFP reporter mice, which faithfully reflect IFNγ protein expression^**35**^. IFNγ production was restricted to CD4 T cells, CD8 T cells, and NK cells (Fig. 5A, S5A), with no IFNγ produced by other immune populations before or after IFNAR blockade during chronic infection (Fig. S5B). IFNAR blockade selectively increased IFNγ production by activated PD-1⁺ CD4 T cells and CD8⁺ T cells, but not NK cells, with PD-1⁺ CD8⁺ T cells exhibiting both increased frequency of IFNγ⁺ cells and elevated per-cell IFNγ expression (Fig. 5A, S5A). Although the proportion of IFNγ-expressing LCMV-GP33 tetramer+ CD8 T cells was unchanged, the total number IFNγ+ LCMV-GP33 tetramer+ CD8 T cells and the per cell expression of IFNγ (gMFI) were increased following IFN-I blockade (Fig. 5B). An analogous increase in the number and single cell expression (gMFI) of IFNγ was also observed by the LCMV-GP276 tetramer+ CD8 T cells (Fig S5C). Further analysis indicated that both the Tpex and Tterm populations, but not the Tint/Teff population, specifically increased in frequency and single cell expression of IFNγ following IFNAR blockade (Fig 5C, S5D). Thus, loss of IFN-I signaling specifically increased the number and amount of IFNγ expression from T cells.

**Figure 5.**
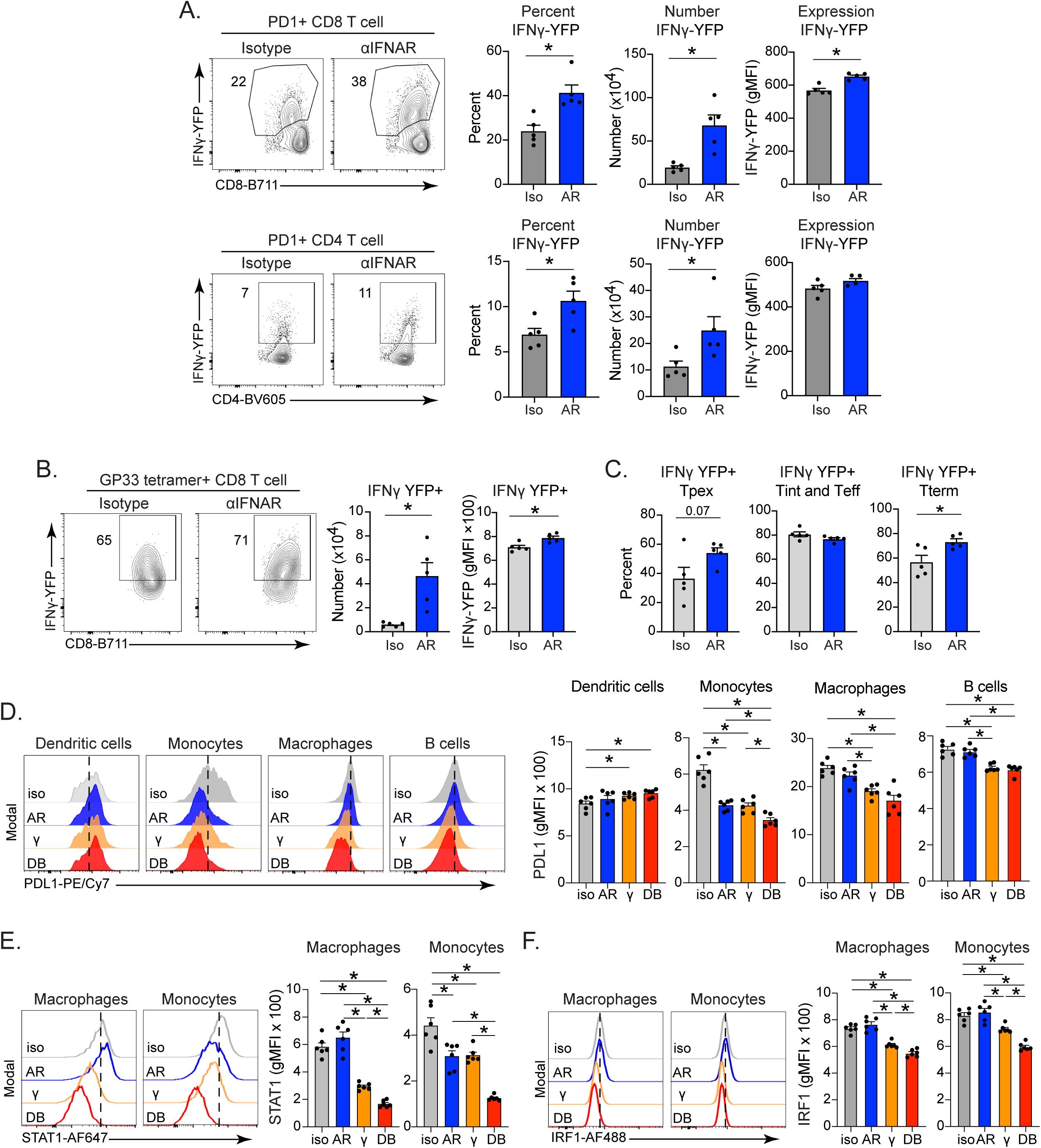
Regulation of IFNγ and PDL1 by IFN-I signaling. **(A)** Flow plots show YFP (IFNγ) expression by splenic PD1+ CD8 T cells (top) and PD1+ CD4 T cells on day 35 of mice treated with isotype (iso) or IFNAR (AR) blocking antibody. Bar graphs show the percent, number, and single-cell expression (gMFI) of YFP (IFNγ) by the PD1+ CD8 T cells (top) and PD1+ CD4 T cells (bottom). **(B)** LCMV-GP33 tetramer+ CD8 T cells after the final isotype (iso) or anti-IFNAR (AR) antibody treatment. Bar graphs show the number and single cell expression (gMFI) of YFP (IFNγ) expressing LCMV-GP33 tetramer+ CD8 T cells. **(C)** Bar graphs show the percent of YFP (IFNγ) expressing Tpex, Tint and Teff combined, and Tterm populations. **(D)** Histograms represent PDL1 expression on day 35 by the indicated cell type in the spleen following isotype (iso), anti-IFNAR (AR), anti-IFNγ (γ), or anti-IFNAR + anti-IFNγ double block (DB). The dotted line in each histogram indicates the average gMFI in the isotype sample. Bar graphs show single-cell expression (gMFI) of PDL1 by the indicted cell type. **(E, F)** Histograms bar graphs represent **(E)** STAT1 and **(F)** IRF1 expression on day 35 by splenic macrophages and monocytes after the indicated blockade. Data are representative of 2 experiments with 5-6 mice per group. The dotted line in each histogram indicates the average gMFI in the isotype sample. Circles in the bar graph indicate individual mice. *, p<0.05 by one-way ANOVA or unpaired t-test.

Because PD-L1 is a key interferon-regulated checkpoint ligand implicated in Tpex suppression¹⁰, we next evaluated its expression across immune populations. PD-L1 expression on macrophages, monocytes, and B cells declined following IFNγ blockade, with combined IFN-I and IFNγ blockade producing the most pronounced reduction (Fig. 5D). In contrast, and unlike the onset of infection^**3,4**^, IFN-I blockade alone had minimal impact on PDL1 expression in most populations, with the exception of monocytes (Fig. 5D). Although modest increases in PDL1 expression on dendritic cells were occasionally observed following antibody blockade, decreases were not detected. Given that both IFN-I and IFNγ regulate PDL1 expression^**36–38**^ and that exhausted T cells closely interact with PDL1 expressing antigen-presenting cells^**17,39,40**^, we hypothesized that elevated IFNγ following IFNAR blockade sustains PD-L1 expression within the chronic inflammatory niche.

Consistent with this model, IFNγR was broadly expressed on DCs, macrophages, monocytes, and B cells and remained unchanged after IFNAR blockade (Fig. S5E). Assessment of downstream signaling revealed that macrophages uniquely maintained elevated STAT1 and IRF1 expression following IFNAR blockade (Fig. 5E, F), indicating sustained IFNγ signaling and correlating with preserved PDL1 expression. Monocytes retained IRF1 with reduced STAT1, suggesting ongoing but attenuated IFNγ signaling relative to macrophages. Although PDL1+ DCs persisted, the greater abundance of macrophages and monocytes in the spleen suggests that these populations principally account for the maintenance of a PDL1-rich splenic environment (Fig. S5F). Further, the IFNγ induced gene MHC II was upregulated on macrophages and monocytes following IFN-I blockade, whereas IFNγ blockade reduced MHC II expression (Fig. S5G). Thus, IFN-I blockade enhances IFNγ production by T cells, which is subsequently sensed by macrophages and monocytes to sustain PDL1 expression and reinforce checkpoint-mediated suppression during chronic infection.

### Interferon-driven PD-L1 enforces Tpex suppression during chronic infection

We next investigated whether reduced PD-L1 expression accounts for the enhanced Tpex expansion observed following combined IFN-I and IFNγ blockade. Virus-specific CD8⁺ T cell responses were therefore compared across conditions of IFNAR plus IFNγ blockade with or without additional PD-L1 inhibition (Fig. 6A). If Tpex expansion following interferon double blockade is mediated through diminished PD-L1–dependent suppression, then PD-L1 blockade would be expected to provide little additional expansion beyond the double interferon blockade alone.

**Figure 6.**
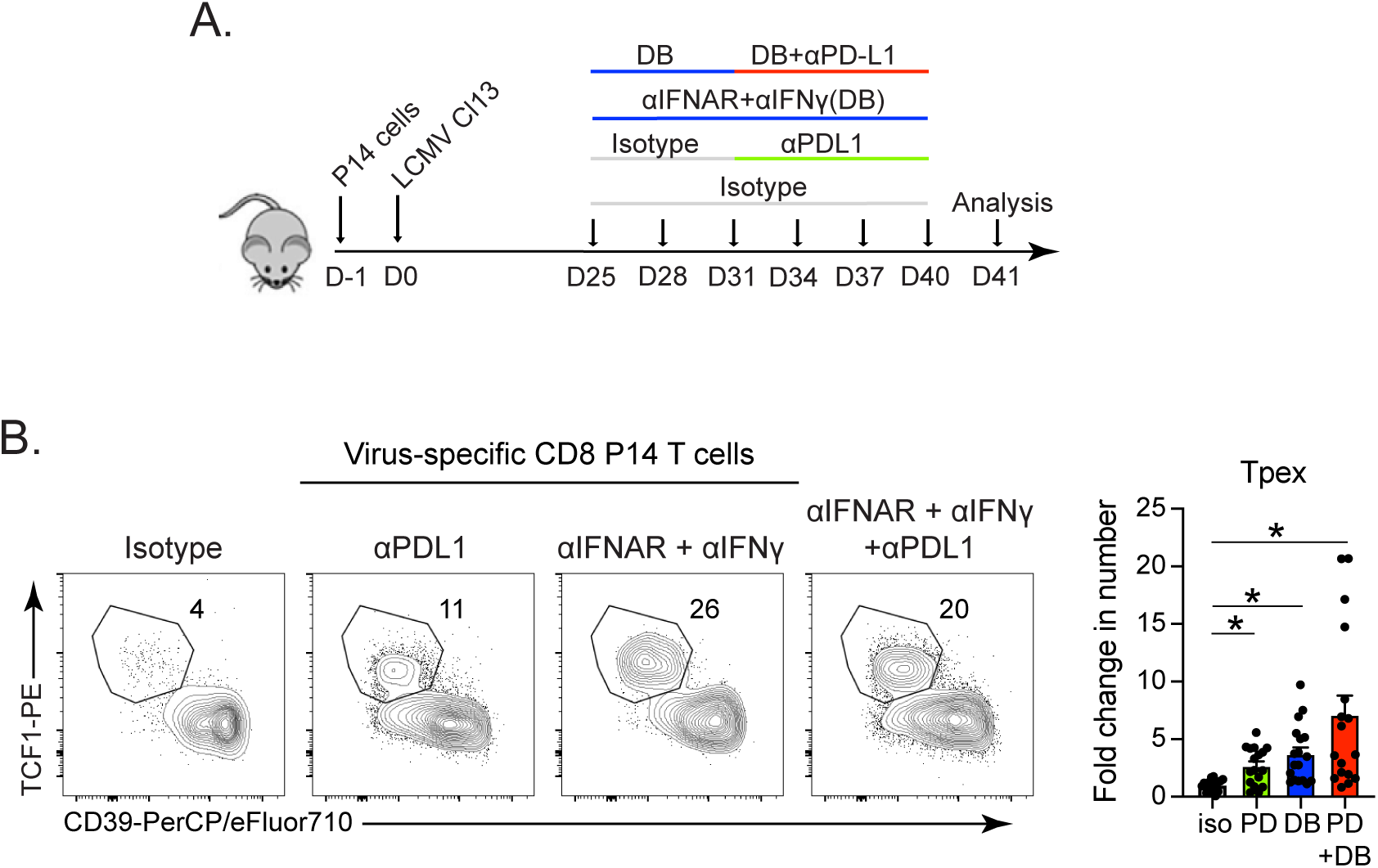
IFN-I and IFNγ modulate PDL1 expression to suppress Tpex proliferation. **(A)** Schematic diagram of the antibody blockade. **(B)** Flow plots show the frequency of Tpex following PDL1 (PD) blockade and anti-IFNAR + anti-IFNγ double block (DB) alone or combined. The bar graph shows the fold change compared to isotype treatment in the number of LCMV-specific CD8 P14 Tpex cells. Data show the compiled results of 3 experiments with 5-6 mice per group. Circles in the bar graph indicate individual mice. *, p<0.05 by one-way ANOVA.

To allow interferon-dependent PDL1 downregulation to occur prior to checkpoint inhibition, mice first received two doses of IFN-blocking antibodies before anti-PDL1 treatment was introduced at the third dose and maintained thereafter (Fig. 6A). An anti–PDL1-only group was included to assess the independent effect of checkpoint blockade, while isotype-treated mice served as controls. Both PDL1 blockade alone and IFNAR + IFNγ double blockade independently increased P14 Tpex numbers to a comparable extent (Fig. 6B). Notably, combining PD-L1 blockade with interferon double blockade did not significantly expand the Tpex compartment beyond either treatment alone (Fig. 6B), indicating that loss of PD-L1 is a major mechanism driving Tpex expansion downstream of IFN-I and IFNγ withdrawal.

## DISCUSSION

Sustained IFN-I exposure during prolonged antigen stimulation, as occurs in chronic viral infection, exerts both activating and suppressive effects on immune cells^**3,4,41**^. However, how interferons regulate CD8 T cell states once exhaustion is firmly established has remained uncertain. Here, we identify a two-layered, interferon-driven regulatory system that controls exhaustion by constraining the expansion and spatial positioning of Tpex during established chronic infection.

Persistent IFN-I signaling limits Tpex proliferation by enforcing their sequestration within B cell follicles, thereby restricting access to proliferative cues. When IFN-I signaling is blocked, Tpex redistribute into the T cell zone of the lymphoid tissue and undergo partial expansion. However, this proliferative response is subsequently curtailed by a second interferon-dependent safe-guard: increased T cell-derived IFNγ sustains PD-L1 expression on macrophages and monocytes, preserving an inhibitory myeloid checkpoint that restrains further Tpex expansion. Mechanistically, neither IFN-I nor IFNγ acts directly on Tpex to suppress proliferation. Instead, interferons minimize proliferation through spatial sequestration and maintenance of myeloid PD-L1 expression. Only combined blockade of IFN-I and IFNγ dismantles both regulatory layers, relieving spatial restriction and checkpoint inhibitions to enable maximal Tpex expansion. Thus, IFN-I enforces a multi-layered regulatory program during chronic infection that simultaneously limits IFNγ production, restrains Tpex expansion, and stabilizes Tex functional integrity, thereby favoring long-term CD8 T cell preservation at the expense of rapid reinvigoration.

During early phase of chronic infection, IFN-Is act as “signal three” to promote CD8 T cell priming and effector differentiation ^**3,4,42,43**^. Consistent with this, IFN-I blockade at infection onset suppresses effector differentiation and biases toward Tpex fates^**13**^, potentially explaining the accelerated viral clearance following early IFN-I blockade^**3,4**^. However, because Tpex exhibit limited cytotoxic capacity compared with Tex^**9,14**^, IFN-I mediated promotion of effector differentiation instead of Tpex formation may be physiologically essential for initial viral control. Indeed, preferential Tpex formation at the expense of effector differentiation compromises control of secondary challenges during chronic disease^**15**^. Thus, by protecting the size and persistence of the Tpex reservoir throughout infection, the interferon system is essential for balancing both immediate and durable viral control.

Mechanistically, IFN-I dependent sequestration of Tpex within B cell regions likely limits antigen exposure and TCR stimulation, as viral burden is reduced in these regions during chronic LCMV infection ^**11,32–34**^. IFN-I blockade permits Tpex redistribution into the T cell zone, where interactions with DCs and macrophages capable of providing TCR and co-stimulatory cues to support proliferation can occur. However, IFN-I withdrawal alone is insufficient to fully relieve suppression. Instead, increased IFNγ production by activated T cells is compensated for by sustaining PDL1 expression on myeloid cells, thereby maintaining an inhibitory checkpoint barrier even after Tpex relocation. Consequently, only combined IFN-I and IFNγ blockade enables redistribution and reduces PDL1 myeloid populations to enable Tpex expansion. Of note, PDL1 blockade has been shown to promote Tpex proliferation through interactions with XCR1+ DCs in the splenic marginal zone^**39**^. However, chronic LCMV infection causes rapid destruction of the marginal zone that never completely reforms^**3,4,44**^, which may explain the altered spatial dynamics following interferon versus PDL1 blockade. By identifying PDL1 as a dominant downstream effector of interferon-mediated Tpex suppression, our findings clarify how cytokine and checkpoint pathways converge to regulate Tpex behavior during established chronic infection.

In addition to these indirect mechanisms, direct IFN-I signaling in exhausted CD8 T cells may serve as a buffering system that restrains excessive activation while preserving effector competence during sustained antigen exposure. Virus-specific CD8 T cells, particularly Tpex, exhibit high IFN-I signaling during chronic infection, which is reduced following IFN-I or combined IFN-I/IFNγ blockade, indicating continuous interferon-dependent tuning of exhausted T cell gene expression. Paradoxically, despite their inflammatory nature, IFN-Is dampen overall inflammatory tone by suppressing IFNγ production and by directly sustaining effector-associated transcriptional programs in both Tpex and Tex populations. The diminution of IFNγ (a cytokine associated with enhanced T cell response^**45**^), while preserving residual effector functions suggests that IFN-I blockade decouples different components in the T cell exhaustion programs. Consistent with this, scRNA-seq analysis revealed broad reduction in transcription factors and signaling intermediates downstream of TCR, NF-κB, and MAPK pathways^**46–48**^ following IFN-I signaling blockade, with more pronounced effects after combined IFN-I and IFNγ inhibition. In Tint (i.e., the immediate Tpex progeny) and terminal effector populations, diminished AP-1, Tbx21, and Bhlhe40 regulon activity, accompanied by increased *Eomes* expression^**27**^, enhanced Tox-associated terminal differentiation programs^**21**^, and a shift toward oxidative and fatty-acid metabolism, would be predicted to accelerate terminal differentiation^**8,49,50**^. Thus, while interferon blockade promotes numerical expansion of the Tpex pool, the absence of their interferon-dependent effector-sustaining signals biases their progeny toward deeper dysfunction.

Collectively, these data reveal a bifurcated role for IFN-I during established chronic infection: interferons indirectly restrain Tpex amplification through spatial organization and myeloid checkpoint regulation, while directly preserving the transcriptional integrity and residual effector capacity of exhausted CD8 T cells. This dual-safeguarded architecture has important implications for immunotherapy. Because checkpoint blockade relies on Tpex as the proliferative reservoir that replenishes effector cells^**10**^, manipulating interferon pathways may reshape both the size and quality of this reservoir. The observation that combined interferon blockade amplifies Tpex, together with the parallels to enhanced clinical responses with combined JAK inhibition and PD-1 blockade in cancer^**51,52**^, suggests that temporally controlled modulation of interferon signaling could enhance checkpoint efficacy. However, because interferons also sustain effector integrity, indiscriminate inhibition risks expanding a dysfunctional pool of exhausted cells that could diminish subsequent therapeutic efficacy, emphasizing the need for precise temporal and combinatorial strategies. Our identification of this interferon-regulated architecture opens new avenues for therapeutically uncoupling Tpex expansion from terminal exhaustion and for optimizing interferon-PDL1-targeted combinations. Moving forward, rational therapeutic strategies will need to precisely uncouple Tpex expansion from terminal exhaustion to restore durable antiviral immunity.

## AUTHOR CONTRIBUTIONS

S.L., D.G.B. designed research. S.L., S.Lukhele, H.J.E., and B.C.B. performed experiments. S.L., R.Q., D.A-R., and D.G.B. analyzed data. W.X., M.Y.R.L, S.Lukhele, S. Lamorte, T.L.M. contributed experimental direction, insight, interpretation, and discussion. S.L. and D.G.B. wrote the paper.

## ACKNOWLEDGEMENTS

We thank past and present members of the Brooks and McGaha laboratories for technical help and discussion. This work was supported by the Canadian Institutes of Health Research (CIHR) Grants FDN148386, PJT505111, PJT 203790 (D.G.B) the National Institutes of Health (NIH) grant AI085043 (D.G.B), and the Scotiabank Research Chair to D.G.B.

## SUPPLEMENTAL FIGURE LEGENDS

**Figure S1.**
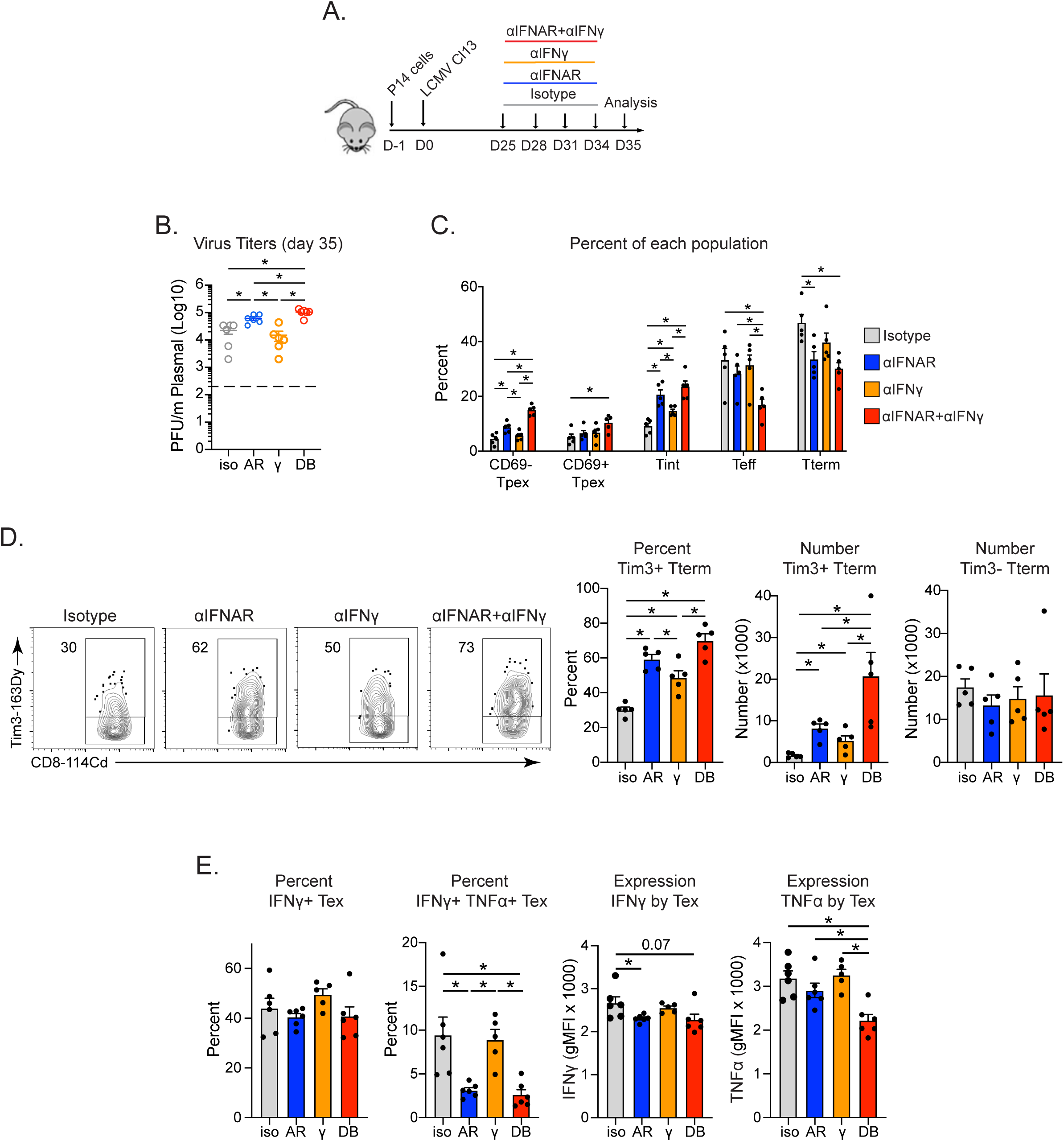
(Related to Figure 1). Virus titers and response of virus-specific CD8 T cells to each antibody blockade. **(A)** Experimental schematic: virus-specific CD8 P14 T cells were transferred one day prior to LCMV-Cl13 infection. Isotype (iso), anti-IFNAR (AR), anti-IFNγ (γ), or combined anti-IFNAR and anti-IFNγ double block (DB) antibody treatment was initiated on day 25 after infection and then every three days following for a total of 4 treatments. Mice were euthanized and analysis was performed one day following the last antibody treatment (on day 35 after infection,) unless otherwise noted. **(B)** Plasma virus titers on day 35, after the last antibody treatment. Dashed line indicates the level of detection of the plaque assay. **(C)** Bars indicate the percent of virus-specific CD8 P14 T cells in each population (defined below the bars) following the indicated antibody treatment. The data are related to the number of these populations in Figure 1E. **(D)** Flow plots show the percent and the bar graph the percent and number of Tim3-positive virus-specific CD8 P14 Tterm cells following each treatment. **(E)** Bar graphs show the percent of IFNγ+, and of IFNγ and TNFα double positive virus-specific CD8 P14 Tex cells, and their single cell (gMFI) expression of IFNγ and TNFα. Data are representative of 3-5 experiments with 5-6 mice per group. Circles indicate individual mice. *, p<0.05 by one-way ANOVA.

**Figure S2.**
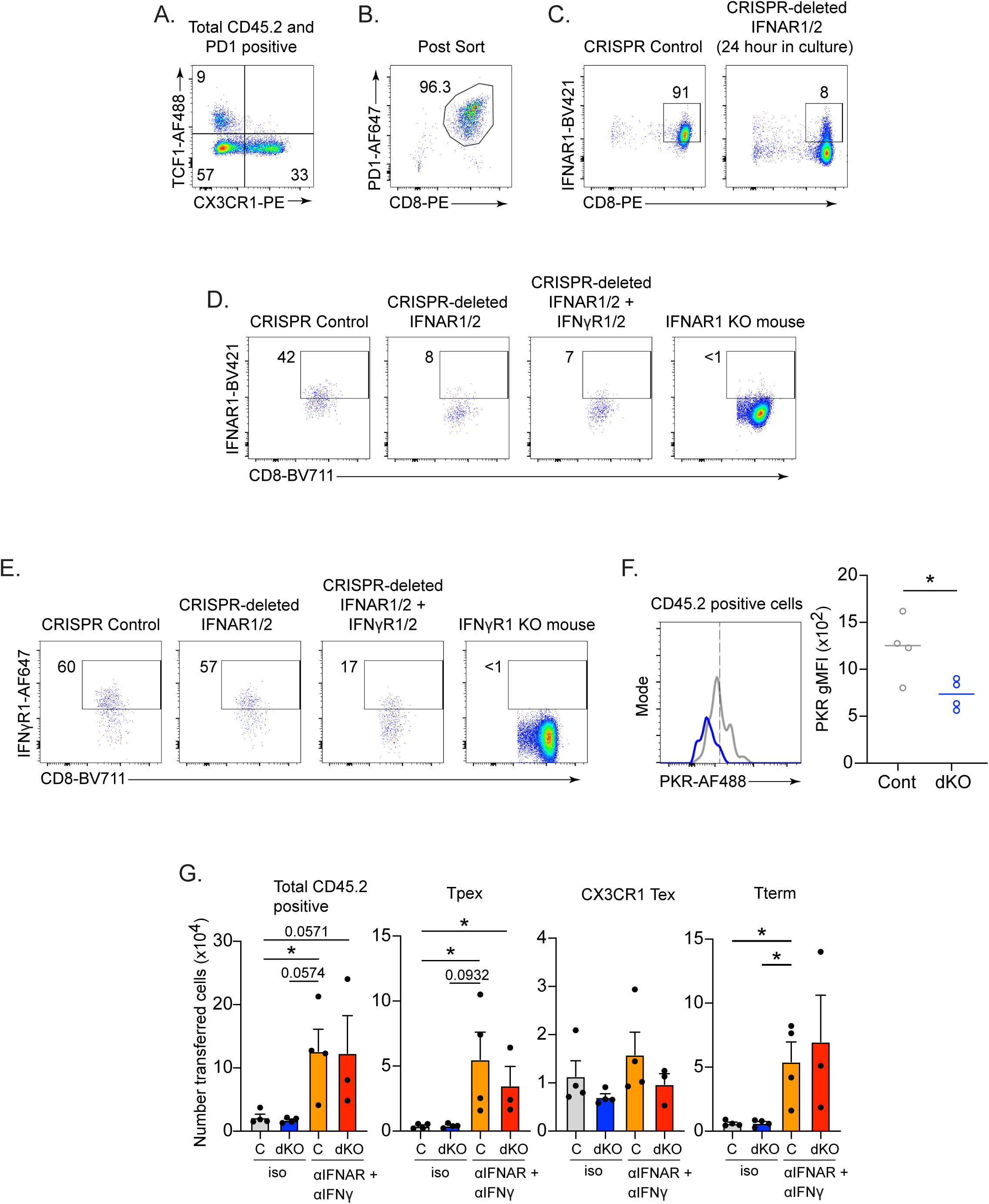
(Related to Figure 2). IFN-I and IFNγ indirectly control Tpex proliferation. **(A, B)** Flow plots show TCF1 and CX3CR1 expression on donor CD45.2+ PD1+ CD8 T cells **(A)** prior to FACSorting (day 25 after infection), and **(B)** post-sort. **(C)** IFNAR1 expression on CRISPR control or IFNAR1 and IFNAR2 CRISPR deleted CD8 T cells 24 hours after CRISPR treatment. **(D, E)** Flow plots show **(D)** IFNAR1 expression and **(E)** IFNγR1 expression on CD45.2+ PD1+ CD8 T cells at the end of the experiment (eight days after transfer). The CRISPR targeting they had received prior to transfer is indicated above each flow plot. IFNAR1 knockout mice and IFNγR1 knockout mice are included as a staining control. **(F)** Histogram shows expression of the type I interferon-stimulated gene PKR in CRISPR control (gray) or dKO (blue) CD45.2+ PD1+ CD8 T cells on day 14 after transfer (day 39 after infection). Bar graph shows the geometric mean fluorescence intensity (gMFI) of PKR staining. **(G)** The experimental design is the same as in the schematic in Figure 1B wherein CRISPR control (C) or double knockout (dKO; IFNAR1, IFNAR2, IFNγR1, and IFNγR2 CRISPR deleted) CD45.2+ PD1+ CD8 T cells were transferred into recipient mice. However, in this experiment, groups of recipient mice were treated with isotype antibody or anti-IFNAR + anti-IFNγ double blockade the day of adoptive transfer. Mice were then euthanized and the indicated populations were enumerated in the spleen 14 days after transfer. Data are representative of 4 experiments, except the data in panel F (treatment with the blocking antibodies with CRISPR deletion), which was only performed one time. 3-5 mice were used per group. Circles in the bar graph and scatter plot indicate individual mice. *, p<0.05 by one-way ANOVA.

**Figure S3.**
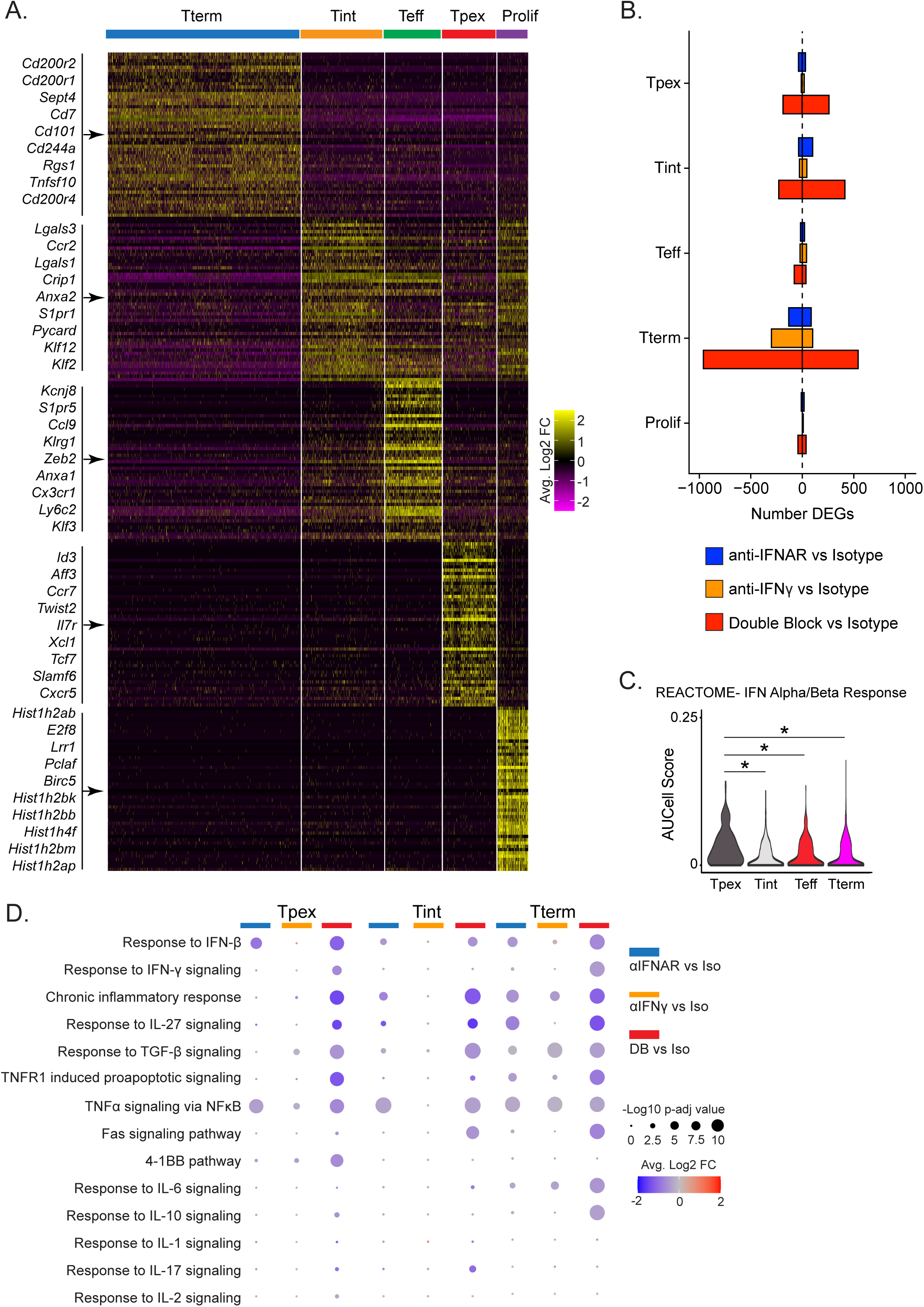
(Related to Figure 3). Transcriptional alterations in virus-specific CD8 T cells following interferon blockade. **(A)** Heatmap shows the top 50 upregulated genes in each cluster, and representative genes for each cluster are provided on the left. **(B)** Bars show the number of DEGs in the indicated cell type in anti-IFNAR (blue), anti-IFNγ (orange), or double blockade (red) versus isotype control. **(C)** Violin plot shows the REACTOME IFN Alpha/Beta Response pathway in Tpex, Tint, Teff, and Tterm cells. *, adj. p-value < 0.05. **(D)** Pathway analysis indicating the Log2 average fold change (FC) by Tpex, Tint, and Tterm following anti-IFNAR, anti-IFNγ, or combined anti-IFNAR and anti-IFNγ double blockade (compared to isotype control treatment).

**Figure S4.**
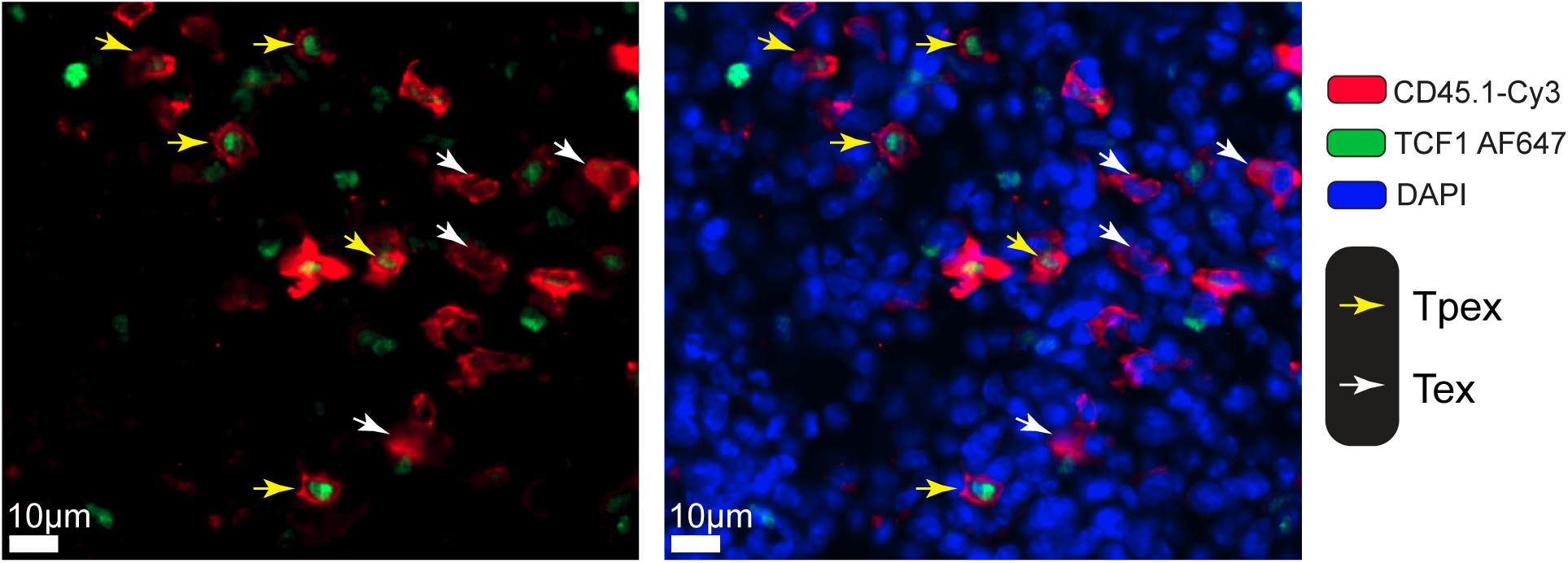
(Related to Figure 4). Tpex and Tex identification. Representative magnified images related to Figure 4 showing Tpex and Tex in the spleen. Yellow arrows indicate Tpex, white arrows indicate Tex. Images are the same, but the left image lacks DAPI to allow better visualization of TCF1 staining.

**Figure S5.**
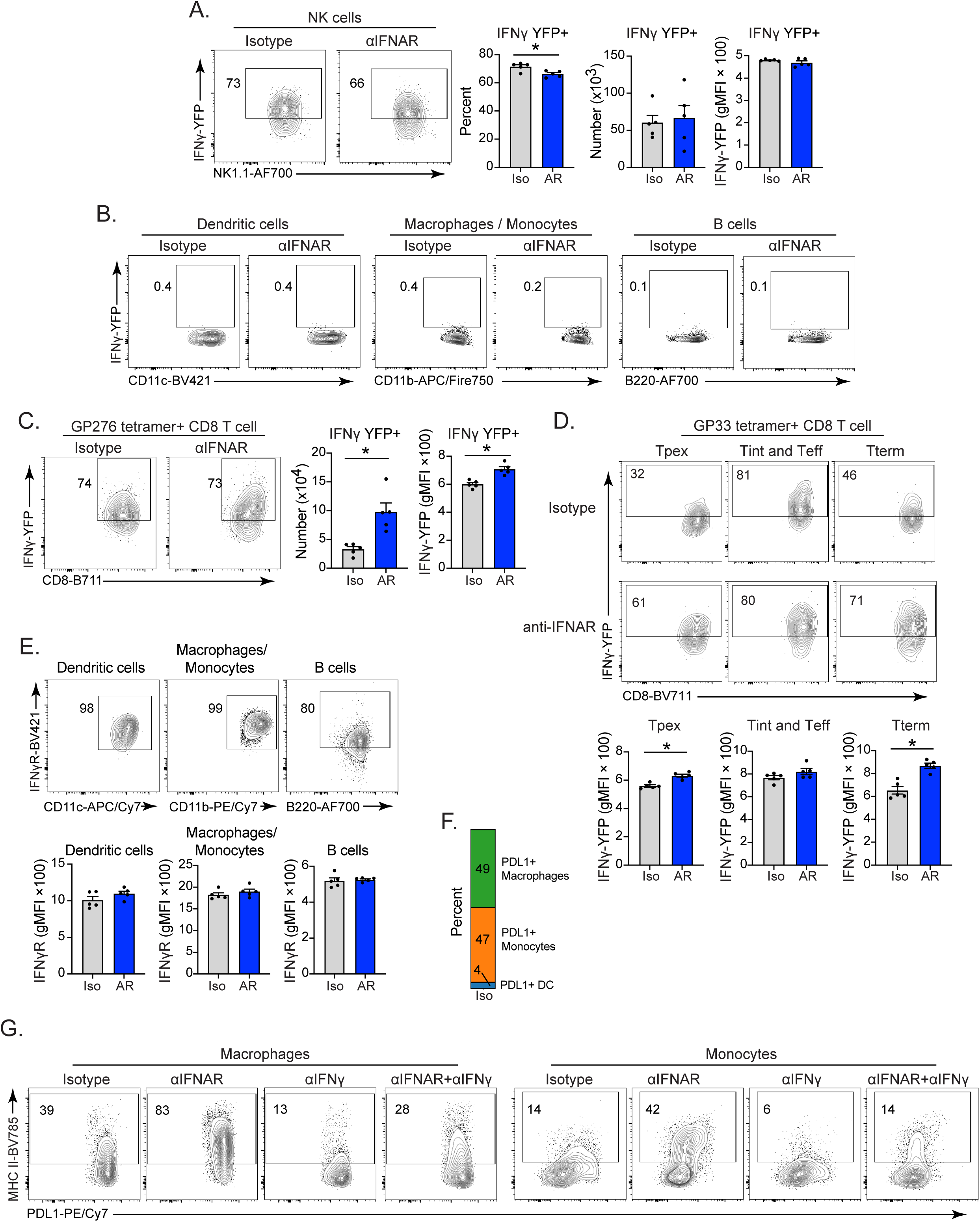
(Related to Figure 5). IFNγ and IFNγR1 expression by population. **(A-B)** Flow plots show YFP (IFNγ) expression by splenic (**A)** NK cells, **(B)** Dendritic cells, macrophages/monocytes and B cells in isotype and anti-IFNAR treated mice. **(C)** LCMV-GP276 tetramer+ CD8 T cells after the final isotype (iso) or anti-IFNAR (AR) antibody treatment. Bar graphs show the number, and single cell expression (gMFI) of YFP (IFNγ) expression in each population. **(D)** Flow plots show YFP (IFNγ) expression by Tpex, Tint and Teff combined, and Tterm populations on the day after the final isotype or anti-IFNAR antibody treatment. Bar graphs show the single cell expression (gMFI) of each population in isotype and anti-IFNAR treated mice. **(E)** IFNγR1 expression by the indicated population on the day after the final antibody treatment. **(F)** The stacked bar graph shows the composition of myeloid populations in the spleen that are PDL1+ macrophages (green), PDL1+ monocytes (orange), and PDL1+ DCs (blue). **(G)** MHC II expression on macrophages and monocytes following the indicated antibody blockade. Data are representative of 2-3 experiments with 5-6 mice per group. The dotted line in each histogram indicates the average gMFI in the isotype sample. Circles in the bar graph indicate individual mice. *, p<0.05 by one-way ANOVA.

## METHODS

### Materials Availability

Requests for resources and reagents should be directed to David Brooks.

#### Mice

C57BL/6 mice were purchased from Jackson Laboratories or the rodent breeding colony at the Princess Margaret Cancer Center. LCMV-GP33-41 specific CD8 P14 TCR transgenic T cells have been described previously^**53**^. IFNAR knockout mice were previously described^**3**^ and generously provided by Dr. Dorian McGavern (National Institutes of Health). IFNγR-/- mice were purchased from Jackson Laboratories (Stock# 003288). *TCF7*-GFP mice were purchased from Jackson Laboratory (Stock# 030909) and crossed with P14 mice. All mice were housed under specific-pathogen free conditions. IFNγ-YFP mice were purchased from Jackson Laboratory (Stock# 017581). Mouse handling conformed to the experimental protocols approved by the Animal Care Committee at the Princess Margaret Cancer Center / University Health Network.

#### Virus Infection and adoptive cell transfer

mice were infected intravenously via the retroorbital sinus with 2×10^6^ PFU of LCMV-Clone13 (Cl13). Viral stocks were prepared and titered by limiting dilution plaque assay as previously described^**54**^. LCMV-specific CD8 P14 transgenic T cells were isolated from the spleens of transgenic mice by negative selection (StemCell Technologies) and transferred i.v. in the retroorbital sinus one to three days prior to infection. Where indicated, C57BL/6 mice received 1000 CD45.1+ LCMV-GP33-41 specific CD8 P14 transgenic T cells. In the microscopy experiments, some mice received 1000 CD45.2+ TCF1-GFP expressing P14 T cells.

#### *In vivo* antibody treatment

WT mice were treated with 500 μg of isotype control antibody, anti-IFNAR and anti-IFNγ blocking antibodies beginning on day 25 and then every 3 days for a total of 4 to 6 treatments, as indicated in Fig. 6 and S1A^**3**^. WT mice received 250 μg anti-PDL1 blocking antibody beginning on day 31 after infection, as indicated in Fig. S6.

### Time-of-Flight mass cytometry (CyTOF)

Up to 4×10^6^ splenocytes were pulsed with 12.5μM Cisplatin (BioVision) in PBS for 1 min prior to quenching with CyTOF staining media (Mg+/Ca+ HBSS containing 2% FBS (Multicell), 10mM HEPES (Cornning), and FBS underlay. Cells were then resuspended in staining media containing metal-tagged surface antibodies (Table S1) and Fc block (CD16/32; in house) for 30 min at 4°C. Cells were fixed, permeabilized and stained with metal tagged intracellular antibodies (Table S1) using the eBioscience™ Foxp3 / Transcription Factor Staining Buffer Set according to manufacturer’s instructions. All antibody concentrations were used at saturating concentrations previously determined by titration. Cells were then incubated overnight in PBS (Multicell) containing 0.3% (ws/v) saponin, 1.6% (v/v) paraformaldehyde (diluted from 16%; Polysciences Inc) and 50 nM Iridium (Fluidigm). Cells were analyzed on a Helios mass cytometer (Fluidigm). EQ Four Element Calibration Beads (Fluidigm) were used to normalize signal intensity over time and data analysis was performed. P14 T cells were gated on (DNA/Iridium+, single event length, cisplatin negative, B220- NK1.1-, CD4-, CD45.2- CD8a+ CD45.1+). UMAP analyses were performed on the P14 cells

### Bioinformatic Analyses (CyTOF)

Data pre-processing and dimensionality reduction of CyTOF data: Preprocessing of files was performed using FlowJo^TM^ Software (v10) software. Samples were manually debarcoded and exported as separated FCS files. P14 cells were filtered by gating on DNA, singlets, live cells, and CD45.1+ CD45.2- CD8+ CD4-, MHCII- cells, and raw signal events were then exported as matrices in csv format. Data was then analyzed in R (v 2022.02.3). All events were included in dimensionality reduction. Marker expression values were arcsinh transformed using a custom co-factor for each marker before clustering. Phenograph and UMAP were performed using the R implementation of the “Rphenograph” package (v 0.99.1) by JinmiaoChen lab on GitHub^**55**^ (https://github.com/JinmiaoChenLab/Rphenograph) and package “umap” (v 0.2.7.0). Differential state and abundance analyses were performed by “diffcyt” package^**56**^ (v 1.14.0).

#### Flow cytometry and intracellular cytokine stimulation

Single cell spleen suspensions were stained *ex vivo* using antibodies to CD4 (GK1.5), CD8 (53-6.7), CD45.1 (A20), CD45.2 (104), CD11c (N418), CD11b (M1/70), NK1.1 (PK136), CD19 (6D5), MHC-II (M5/114.15.2), Ly6C (HK1.4), Ly6G (1A8), B220 (RA3-6B2), SLAMF6 (330-AJ), TCF1 (S33-966), CD39 (24DMS1), Granzyme B (QA16A02), PD1 (29F.1A12), PDL1 (10F9G2), CXCR5 (L138D7), IFNγ (XMG1.2), TNFα (MP6-XP22), IFNγR (2E2), IFNAR1 (MAR1-5A3), TOX (NAN448B), CD28(E18), CD69 (H1.2F3), CX3CR1 (SA011F11), STAT1 (1/Stat1), IRF1 (D5E4), Zbtb46 (U4-1374) PKR (epr19374), Ki67 (SolA15). Ki67 and CD39 was purchased from Thermofisher, PKR was purchased from Abcam, TCF1, Zbtb46, TOX, STAT1 were purchased from BD Biosciences, IRF1 was purchased from Cell Signaling Technology, all other antibodies were purchased from Biolegend. LCMV-GP_33-41_ and LCMV-GP_276-285_ MHC-I tetramers were provided by the NIH Tetramer Core Facility. Live/dead staining was done using zombie aqua (Biolegend). Staining was performed as directed using the Foxp3 Transcription Factor Staining kit (Thermofisher). Samples were run on a FACS Lyrics (BD Biosciences) and data analyzed using FlowJo software (BD biosciences).

CD8 P14 T cells were CD4- CD8+ CD45.1+ and endogenous GP33-41 or GP276-286 specific CD8 T cells were defined based on CD4- CD8+ CD45.2+ and GP33-41 tetramer+ or GP276-286 tetramer+. CD8 P14 or tetramer+ Tpex were further defined as TCF1+ and/or SLAMF6+, CD69+/-, CX3CR1- cells. CD8 Tint cells were further defined as TCF1- and/or SLAMF6-, CX3CR1+, Ki67+, CD69-. CD8 Teff cells were further defined as TCF1- and/or SLAMF6-, CX3CR1+, Ki67-, CD69-cells. CD8 Tterm were defined as TCF1- and/or SLAMF6-, CD39+, CD69+ cells. Dendritic cells are defined as CD19-, NK1.1-, Ly6G-, CD3-, Ly6C-, Zbtb46+, CD11c high, MHC-II high. Macrophages are defined as CD19-, NK1.1-, Ly6G-, CD3-, Ly6C-, CD11b+, CD11c intd. Monocytes are defined as CD19- NK1.1-, Ly6G-, CD3-, CD11b+ Ly6C+. B cells are defined as CD19+, B220+, MHC-II+.

For cytokine quantification, splenocytes were restimulated for 5 hours at 37°C with 2μg/ml of the MHC class I-restricted LCMV peptide GP_33-41_ in the presence of 50 U/ml recombinant murine IL-2 and 1 mg/ml brefeldin A (Sigma). Following the 5 hours *in vitro* restimulation, cells were stained with a fixable viability stain, zombie aqua (Biolegend), extracellularly stained as above with CD8, and fixed, permeabilized (Biolegend cytokine staining buffer) and stained with IFNγ (XMG1.2) and TNFα (MP6-XT22). Samples were run on a FACS Lyrics (BD Biosciences) and data analyzed using FlowJo software.

#### CRISPR mediated gene deletion

PD1+ CD8+ T cells were FACSorted from spleens of LCMV-Cl13-infected mice (day 25 p.i.) and resuspended in 20 μl buffer P3 (P3 Primary Cell 4D-Nucleofector X Kit S, Lonza, cat. no. V4XP-3032). 5×10^6^ to 1×10^7^ of cells were used per nucleofection. Single guide RNAs (sgRNAs) were incubated together with recombinant Cas9 (TrueCut Cas9 Protein v2, 5ug/ul, Thermo Fisher Scientific, cat. no. A36499) for 15 min at room temperature, at a ratio of 1:3.3 (i.e., 30 pmol Cas9 protein per 100 pmol sgRNA) to form the CRISPR–Cas9–sgRNA-ribonucleoprotein (RNP) complex. To increase deletion efficiency, a combination of three for each different pre-validated sgRNAs against IFNAR1 + IFNAR2 or IFNγR1 + IFNγR2 were mixed with the cell suspension (see list below) and transferred into a 16-well nucleocuvette strip (Lonza, cat. no. V4XP-3032). For negative control, sgRNA (TrueGuide™ sgRNA Negative Control, non-targeting 1; Thermo Fisher Scientific, cat. no. A35526t) was annealed with Cas9 at the same molar ratio. Cells were nucleofected using program DS137 and buffer P3 on the 4D-Nucleofector system (4D-Nucleofector X unit, Lonza, cat. no. AAF-1003X). Cells were then washed and transferred intravenously via the retroorbital sinus into LCMV-Cl13 infection time-matched mice (100k to 150k cells per mice). The knockout efficiency based on IFNAR or IFNγR expression was confirmed by flow cytometry.

#### sgRNA sequences against mouse IFNAR1/2, IFNγR1/2-

gRNAs were purchased from IDT Mm.Cas9.IFNAR1.1.AA Mm.Cas9.IFNAR1.1.AB

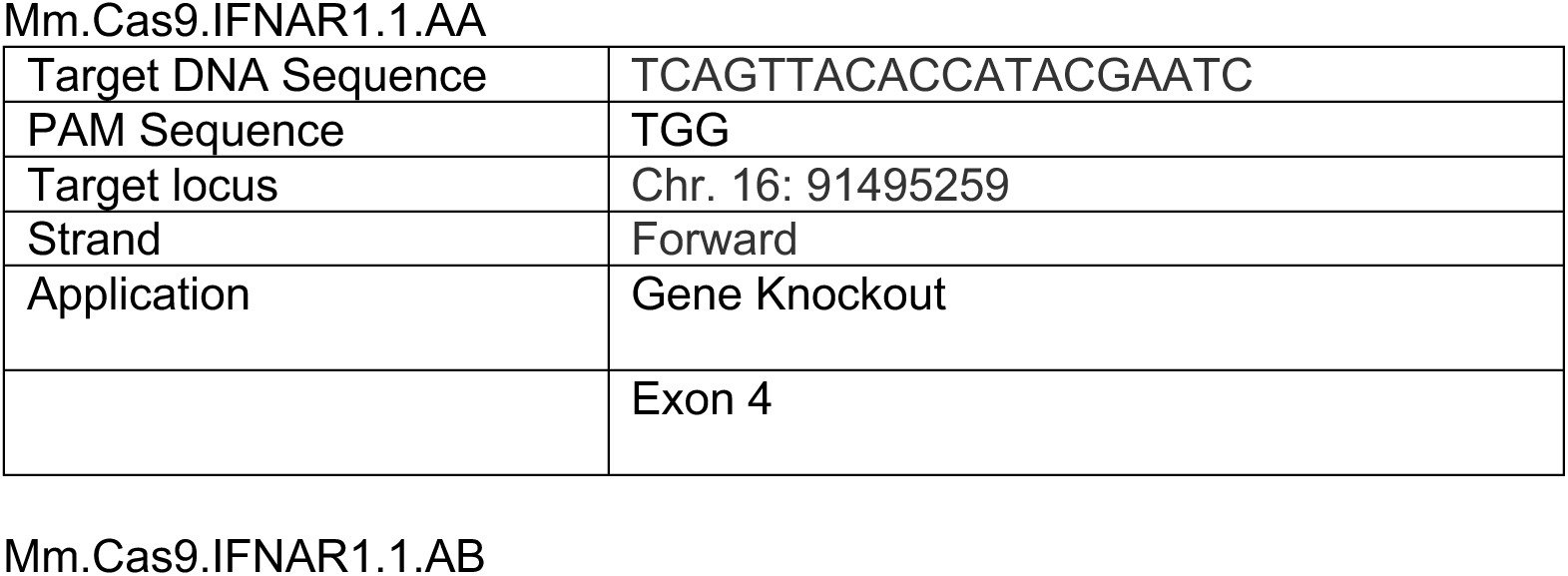
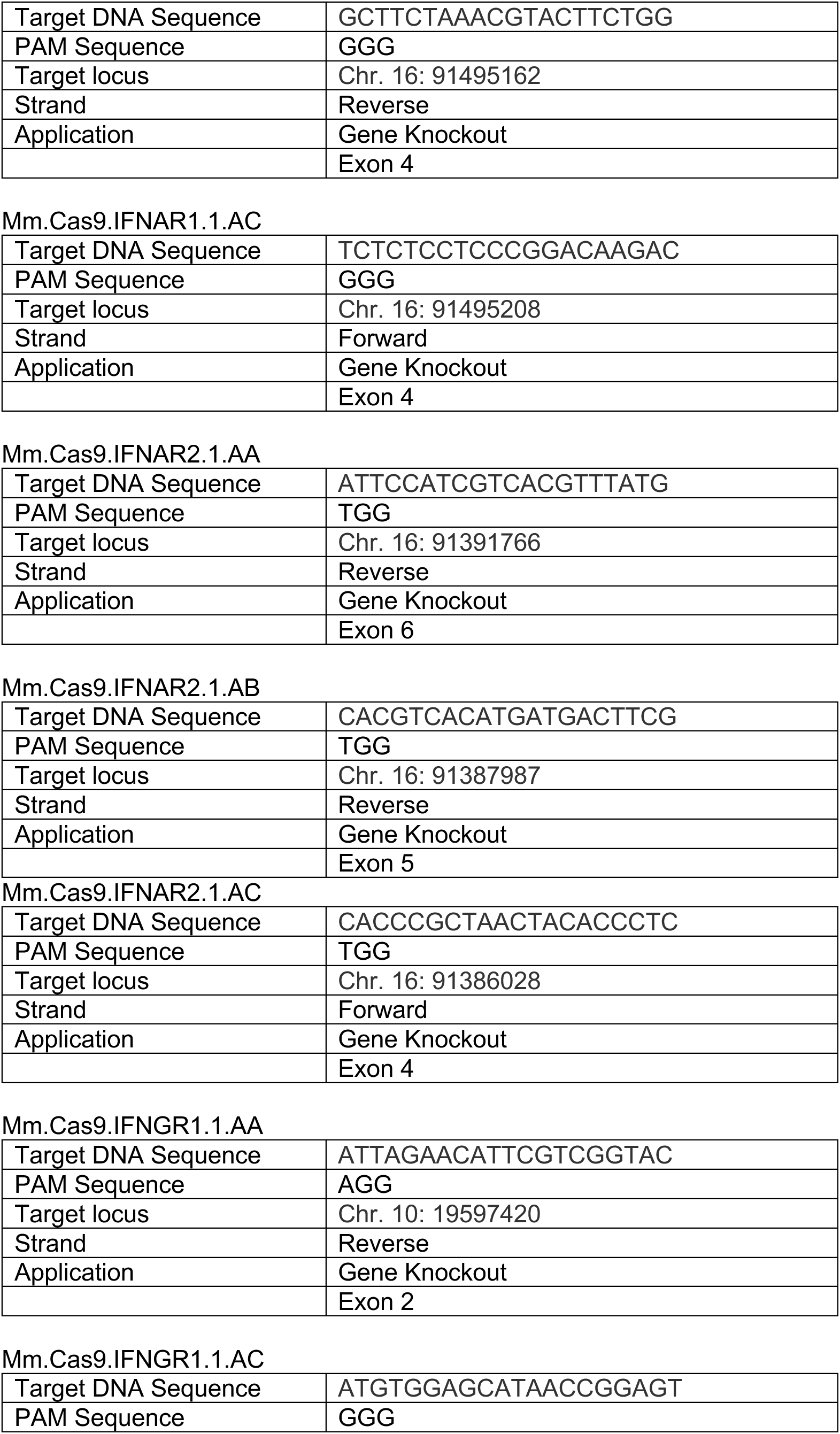
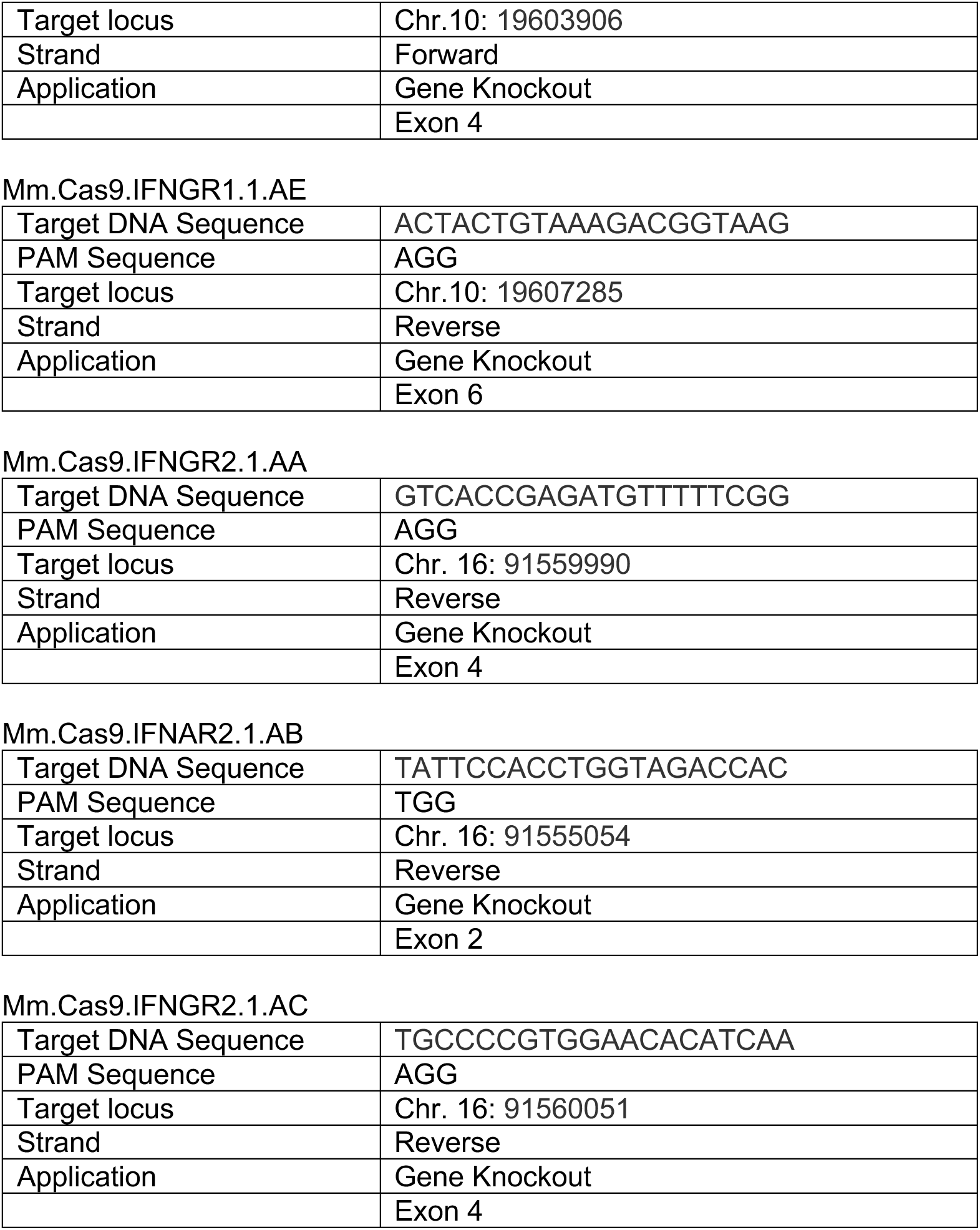

#### Single Cell (sc) RNA sequencing

CD8 P14 T cells were transferred into WT mice one day prior to infection. Mice were treated with anti-IFNAR and/or IFNγ blocking antibodies as shown in Fig S1A. Mice were sacrificed one day after the last treatment (day 33 after infection) and spleens were harvested, digested into single cell suspension, and pooled within each condition. Spleens were pooled from 5-6 mice per group. CD8 T cells were enriched on autoMACS Pro Separator (Miltenyi Biotec) with CD8 positive selection beads (Miltenyi Biotec), and then FACSorted on a BD FACSaria Fusion cell sorter for P14 cells based on CD45.1 and CD8 expression. After sorting and purity check on flow cytometer, P14 cells were then resuspended in 10X Genomics Chromium single cell RNA master mix and loaded onto a 10x chromium chip. The cDNA Library was generated using the Chromium Single Cell 3’ Reagents Kits v2 User Guide:CG00052 Rev B and then sequenced on the NovaSeq X platform to achieve an average of 50,000 reads per cell. Sequencing was performed at the Princess Margaret Genomics Center.

### Bioinformatic Analyses (scRNA-Seq)

#### Reads alignment and quantification

Base calling was performed using Illumina RTA (v 3.4.4) and bcl2fastq2 (v 2.20) to generate bcl files. Cell ranger (v 7.0.0) was then used to demultiplex bcl into fastq files and to align reads to the mm10 genome (refdata-gex-mm10-2020-A downloaded from the 10x genomics website).

#### Doublets identification

Single-cell data was analysed in R (v 4.2.2) and the R Seurat (v 4.9.9.9059) package (Hao et al. 2021)[]. Initial clustering for each sample was performed following the standard Seurat workflow (i.e. NormalizeData, FindVariableFeatures, ScaleData, RunPCA, FindNeighbors, and FindClusters) with default parameters and a resolution of 1. Doublet scores were computed by the cxds co-expression approach from the scds package using the top 3000 highly variable genes^**57**^. Doublet scores were standardized to Z-scores; cells with Z-score > 2 were flagged as potential doublets. Clusters containing > 10% potential doublets were identified as “doublet clusters” and were each subjected re-clustering. To identify sub-clusters that are enriched for doublets, we calculated the delta between individual cell Z-scores and the sample median Z-score. Sub-clusters with a modified Z-score > 4 (derived from the median of these deltas) were classified as doublets and excluded from downstream analyses.

#### Quality control

Cells considered dead or of poor quality were identified as outliers based on the Median Absolute Deviation (MAD) using the isOutlier function from the scuttle package. Specifically, dead cells were defined as outliers based on mitochondrial gene percentage (settings: nmads=5, type=”higher”, min.diff=12) or if they expressed less than 200 genes. Poor-quality cells were identified as outliers for both RNA read counts (nmads=300, type=”higher”, log=T) and feature counts (nmads=400, type=”higher”, log=T). Furthermore, cells were excluded based on the additional following criteria: ribosomal gene percentage > 5%, hemoglobin percentage < 0.025, and a log10GenesPerUMI > 0.8.

The number of cells remaining in each sample following these filters is as follows:

- P14 Isotype: 2440 cells
- P14 BA: 1847 cells
- P14 BG: 1507 cells
- P14 DB: 2417 cells

Genes expressed in less than 10 cells were excluded from further analyses.

#### Sample integration

Individual samples were preprocessed following the standard Seurat workflow (i.e. NormalizeData, FindVariableFeatures, ScaleData, RunPCA). Samples within each group were integrated using the IntegrateLayers function with the Harmony algorithm^**58**^. Nearest-neighbors graph construction was performed using the first 40 principle components from the Harmony-integrated reduction. Finally, cells from P14 and SMT were clustered using resolution 0.2 and 0.1, respectively.

#### Differential gene expression

Differential gene expression analysis was conducted using the Wilcoxon rank-sum test implemented in Seurat, with Bonferroni correction applied for multiple hypothesis testing across predefined cell populations. Genes were considered significantly differentially expressed, if they exhibited an absolute log2 fold change > 0.25 and were expressed in at least 10% of cells in one of the compared groups.

#### Pathway and Regulon analysis

Pathway activity scores were inferred using AUCell^**29**^, applied to gene sets from the msigdb^**59**^ and GO databases. Mus_musculus database. AUCell scores were calculated for the mh, m2, and m5 collections to assess relative pathway activity at the single-cell level. Regulon activity was inferred using SCENIC, following standard preprocessing and scoring procedures to quantify transcription factor regulon activity across cells.

For pathway activity and regulon scores, differential analyses between groups were also performed using the Wilcoxon rank-sum test. To quantify effect size, Cohen’s d was used in place of the default log2 fold-change metric, providing a standardized measure of differences between conditions.

##### Immunofluorescent microscopy and Image analysis

Fresh spleens were fixed in fixation buffer (4%PFA+5%Sucrose+PBS) overnight at 4 °C with gentle shaking. Fixed spleens were then transferred to 10% Sucrose buffer for 1 h, followed by transfer to 30% Sucrose buffer for dehydration overnight at 4 °C with gentle shaking. Fixed spleens were then embedded and frozen by using OCT on dry ice, then stored at -80 °C. Tissue was sliced at 6um by Leica Cyrostat (CM3050S) and mounted on Fisher Scientific Epredia Superfrost Plus slides. Procedures for staining: spleen sections were blocked by using biotin/avidin blockade kit (Vector Laboratories, SP-2001), followed by staining blockade buffer (5% Donkey Serum+0.2% TritonX 100+ PBS) for 1h at room temperature. Primary antibodies: B220-AF488(RA3-6B2, Biolgend), CD4-biotin(RM4-5, Biolegend), Goat-anti-PD-L1(Polyclonal, Cat. AF1019, R&D System); CD4-AF488(RM4-5, Biolegend), CD45.1-biotin(A20, Biolgend), rabbit-anti-TCF1(C63D9, Cell Signaling Technology), B220-AF750(RA3-6B2, R&D System); chicken-anti-GFP(Polyclonal, ab13970, Abcam), CD45.2-biotin(104, Biolegend), CD4-AF647(RM4-5, Biolegend); stained overnight at 4 °C. Secondary antibodies: streptavidin-Cy3(Biolegend), donkey-anti-goat AF647(ab150135, Abcam), donkey anti-chicken AF488(A78948, Thermofisher Scientific), donkey anti-rabbit AF647(A-31573, Thermofisher Scientific), stained for 1h at room temperature. DAPI was stained at 2.5 × 10^-3^ mg/mL for 5 min at room temperature. Tissue slices were mounted by using Thermo Prolong Glass Antifade mountant (P36980) by using #1.5 coverglass (Fisherbrand 12541024CA). Images were acquired at 20X magnification by using Zeiss AxioImager and Scanner Tpex were identified as CD45.1+, TCF1+, DAPI+ cells, or CD45.2+, GFP+, DAPI+ cells. The B cell region is defined by the positive staining of B220; the inner and outer borders of the B cell region are defined by the sharp contrast of B220-positive stain in the B cell region and negative stain in the T cell region and in the red pulp. Tpex located in this area are considered as Tpex in the B cell region. The T cell region is defined by the positive staining of CD4 surrounded by the B220 stain. It should be noted that CD4 T cells infiltrate into the B cell region at the T-B cell boundary, resulting in a CD4 and B220 overlapping region. For our study, this area is defined as B cell region. Only Tpex located within CD4 positive staining regions and located at least 1 cell distance from the edge of the B cell border are quantified as Tpex in the T cell region. Remaining area that is negative for both CD4 and B220 is the red pulp. Images were analyzed in QuPath (v 0.5.1) and HALO 4.0 Software (Indica Lab).

### Statistical analyses (other than CyTOF and scRNAseq)

Student’s T-test or one-way ANOVA were calculated by GraphPad Prism 9 (GraphPad Software, Inc.) as indicated in the Figure legends.

## REFERENCES

1. Lukhele, S., Boukhaled, G.M., and Brooks, D.G. (2019). Type I interferon signaling, regulation and gene stimulation in chronic virus infection. Semin Immunol 43, 101277. 10.1016/j.smim.2019.05.001.

2. Wilson, E.B., and Brooks, D.G. (2013). Decoding the complexity of type I interferon to treat persistent viral infections. Trends Microbiol 21, 634–640. 10.1016/j.tim.2013.10.003.

3. Wilson, E.B., Yamada, D.H., Elsaesser, H., Herskovitz, J., Deng, J., Cheng, G., Aronow, B.J., Karp, C.L., and Brooks, D.G. (2013). Blockade of chronic type I interferon signaling to control persistent LCMV infection. Science 340, 202–207.

4. Teijaro, J.R., Ng, C., Lee, A.M., Sullivan, B.M., Sheehan, K.C., Welch, M., Schreiber, R.D., de la Torre, J.C., and Oldstone, M.B. (2013). Persistent LCMV infection is controlled by blockade of type I interferon signaling. Science 340, 207–211.

5. Wherry, E.J., and Kurachi, M. (2015). Molecular and cellular insights into T cell exhaustion. Nat Rev Immunol 15, 486–499. 10.1038/nri3862.

6. Angelosanto, J.M., Blackburn, S.D., Crawford, A., and Wherry, E.J. (2012). Progressive loss of memory T cell potential and commitment to exhaustion during chronic viral infection. J Virol 86, 8161–8170. 10.1128/JVI.00889-12.

7. Beltra, J.C., Manne, S., Abdel-Hakeem, M.S., Kurachi, M., Giles, J.R., Chen, Z., Casella, V., Ngiow, S.F., Khan, O., Huang, Y.J., et al. (2020). Developmental Relationships of Four Exhausted CD8(+) T Cell Subsets Reveals Underlying Transcriptional and Epigenetic Landscape Control Mechanisms. Immunity 52, 825–841 e828. 10.1016/j.immuni.2020.04.014.

8. Chen, W., Teo, J.M.N., Yau, S.W., Wong, M.Y., Lok, C.N., Che, C.M., Javed, A., Huang, Y., Ma, S., and Ling, G.S. (2022). Chronic type I interferon signaling promotes lipid-peroxidation-driven terminal CD8(+) T cell exhaustion and curtails anti-PD-1 efficacy. Cell Rep 41, 111647. 10.1016/j.celrep.2022.111647.

9. Utzschneider, D.T., Charmoy, M., Chennupati, V., Pousse, L., Ferreira, D.P., Calderon-Copete, S., Danilo, M., Alfei, F., Hofmann, M., Wieland, D., et al. (2016). T Cell Factor 1-Expressing Memory-like CD8(+) T Cells Sustain the Immune Response to Chronic Viral Infections. Immunity 45, 415–427. 10.1016/j.immuni.2016.07.021.

10. Im, S.J., Hashimoto, M., Gerner, M.Y., Lee, J., Kissick, H.T., Burger, M.C., Shan, Q., Hale, J.S., Lee, J., Nasti, T.H., et al. (2016). Defining CD8+ T cells that provide the proliferative burst after PD-1 therapy. Nature 537, 417–421. 10.1038/nature19330.

11. He, R., Hou, S., Liu, C., Zhang, A., Bai, Q., Han, M., Yang, Y., Wei, G., Shen, T., Yang, X., et al. (2016). Follicular CXCR5- expressing CD8(+) T cells curtail chronic viral infection. Nature 537, 412–428. 10.1038/nature19317.

12. Kallies, A., Zehn, D., and Utzschneider, D.T. (2020). Precursor exhausted T cells: key to successful immunotherapy? Nat Rev Immunol 20, 128–136. 10.1038/s41577-019-0223-7.

13. Wu, T., Ji, Y., Moseman, E.A., Xu, H.C., Manglani, M., Kirby, M., Anderson, S.M., Handon, R., Kenyon, E., Elkahloun, A., et al. (2016). The TCF1-Bcl6 axis counteracts type I interferon to repress exhaustion and maintain T cell stemness. Sci Immunol 1. 10.1126/sciimmunol.aai8593.

14. Giles, J.R., Ngiow, S.F., Manne, S., Baxter, A.E., Khan, O., Wang, P., Staupe, R., Abdel-Hakeem, M.S., Huang, H., Mathew, D., et al. (2022). Shared and distinct biological circuits in effector, memory and exhausted CD8(+) T cells revealed by temporal single-cell transcriptomics and epigenetics. Nat Immunol 23, 1600–1613. 10.1038/s41590-022-01338-4.

15. Snell, L.M., MacLeod, B.L., Law, J.C., Osokine, I., Elsaesser, H.J., Hezaveh, K., Dickson, R.J., Gavin, M.A., Guidos, C.J., McGaha, T.L., and Brooks, D.G. (2018). CD8(+) T Cell Priming in Established Chronic Viral Infection Preferentially Directs Differentiation of Memory-like Cells for Sustained Immunity. Immunity 49, 678–694 e675. 10.1016/j.immuni.2018.08.002.

16. Osokine, I., Snell, L.M., Cunningham, C.R., Yamada, D.H., Wilson, E.B., Elsaesser, H.J., de la Torre, J.C., and Brooks, D. (2014). Type I interferon suppresses de novo virus-specific CD4 Th1 immunity during an established persistent viral infection. Proc Natl Acad Sci U S A 111, 7409–7414. 10.1073/pnas.1401662111.

17. Humblin, E., Korpas, I., Lu, J., Filipescu, D., van der Heide, V., Goldstein, S., Vaidya, A., Soares-Schanoski, A., Casati, B., Selvan, M.E., et al. (2023). Sustained CD28 costimulation is required for self-renewal and differentiation of TCF-1(+) PD-1(+) CD8 T cells. Sci Immunol 8, eadg0878. 10.1126/sciimmunol.adg0878.

18. Beltra, J.C., Abdel-Hakeem, M.S., Manne, S., Zhang, Z., Huang, H., Kurachi, M., Su, L., Picton, L., Ngiow, S.F., Muroyama, Y., et al. (2023). Stat5 opposes the transcription factor Tox and rewires exhausted CD8(+) T cells toward durable effector-like states during chronic antigen exposure. Immunity 56, 2699–2718 e2611. 10.1016/j.immuni.2023.11.005.

19. Chen, Z., Ji, Z., Ngiow, S.F., Manne, S., Cai, Z., Huang, A.C., Johnson, J., Staupe, R.P., Bengsch, B., Xu, C., et al. (2019). TCF-1-Centered Transcriptional Network Drives an Effector versus Exhausted CD8 T Cell-Fate Decision. Immunity 51, 840–855 e845. 10.1016/j.immuni.2019.09.013.

20. Kasmani, M.Y., Zander, R., Chung, H.K., Chen, Y., Khatun, A., Damo, M., Topchyan, P., Johnson, K.E., Levashova, D., Burns, R., et al. (2023). Clonal lineage tracing reveals mechanisms skewing CD8+ T cell fate decisions in chronic infection. J Exp Med 220. 10.1084/jem.20220679.

21. Seo, H., Chen, J., Gonzalez-Avalos, E., Samaniego-Castruita, D., Das, A., Wang, Y.H., Lopez-Moyado, I.F., Georges, R.O., Zhang, W., Onodera, A., et al. (2019). TOX and TOX2 transcription factors cooperate with NR4A transcription factors to impose CD8(+) T cell exhaustion. Proc Natl Acad Sci U S A 116, 12410–12415. 10.1073/pnas.1905675116.

22. Connolly, K.A., Kuchroo, M., Venkat, A., Khatun, A., Wang, J., William, I., Hornick, N.I., Fitzgerald, B.L., Damo, M., Kasmani, M.Y., et al. (2021). A reservoir of stem-like CD8(+) T cells in the tumor-draining lymph node preserves the ongoing antitumor immune response. Sci Immunol 6, eabg7836. 10.1126/sciimmunol.abg7836.

23. Vardhana, S.A., Hwee, M.A., Berisa, M., Wells, D.K., Yost, K.E., King, B., Smith, M., Herrera, P.S., Chang, H.Y., Satpathy, A.T., et al. (2020). Impaired mitochondrial oxidative phosphorylation limits the self-renewal of T cells exposed to persistent antigen. Nat Immunol 21, 1022–1033. 10.1038/s41590-020-0725-2.

24. Tang, Y., Chen, Z., Zuo, Q., and Kang, Y. (2024). Regulation of CD8+ T cells by lipid metabolism in cancer progression. Cell Mol Immunol 21, 1215–1230. 10.1038/s41423-024-01224-z.

25. Chen, Y., Zander, R.A., Wu, X., Schauder, D.M., Kasmani, M.Y., Shen, J., Zheng, S., Burns, R., Taparowsky, E.J., and Cui, W. (2021). BATF regulates progenitor to cytolytic effector CD8(+) T cell transition during chronic viral infection. Nat Immunol 22, 996–1007. 10.1038/s41590-021-00965-7.

26. Salmon, A.J., Shavkunov, A.S., Miao, Q., Jarjour, N.N., Keshari, S., Esaulova, E., Williams, C.D., Ward, J.P., Highsmith, A.M., Pineda, J.E., et al. (2022). BHLHE40 Regulates the T-Cell Effector Function Required for Tumor Microenvironment Remodeling and Immune Checkpoint Therapy Efficacy. Cancer Immunol Res 10, 597–611. 10.1158/2326-6066.CIR-21-0129.

27. Paley, M.A., Kroy, D.C., Odorizzi, P.M., Johnnidis, J.B., Dolfi, D.V., Barnett, B.E., Bikoff, E.K., Robertson, E.J., Lauer, G.M., Reiner, S.L., and Wherry, E.J. (2012). Progenitor and terminal subsets of CD8+ T cells cooperate to contain chronic viral infection. Science 338, 1220–1225. 10.1126/science.1229620.

28. Yin, P., Yang, J., Jiang, Y., Han, L., Xiong, T., Liu, H., Fang, Y., Ruan, W., Wu, J., Chen, L., et al. (2025). Enhanced FOS expression improves tumor clearance and resists exhaustion in NR4A3-deficient CAR T cells under chronic antigen exposure. Sci Adv 11, eadw3571. 10.1126/sciadv.adw3571.

29. Aibar, S., Gonzalez-Blas, C.B., Moerman, T., Huynh-Thu, V.A., Imrichova, H., Hulselmans, G., Rambow, F., Marine, J.C., Geurts, P., Aerts, J., et al. (2017). SCENIC: single-cell regulatory network inference and clustering. Nat Methods 14, 1083–1086. 10.1038/nmeth.4463.

30. Lukhele, S., Rabbo, D.A., Guo, M., Shen, J., Elsaesser, H.J., Quevedo, R., Carew, M., Gadalla, R., Snell, L.M., Mahesh, L., et al. (2022). The transcription factor IRF2 drives interferon-mediated CD8(+) T cell exhaustion to restrict anti-tumor immunity. Immunity 55, 2369–2385 e2310. 10.1016/j.immuni.2022.10.020.

31. Man, K., Gabriel, S.S., Liao, Y., Gloury, R., Preston, S., Henstridge, D.C., Pellegrini, M., Zehn, D., Berberich-Siebelt, F., Febbraio, M.A., et al. (2017). Transcription Factor IRF4 Promotes CD8(+) T Cell Exhaustion and Limits the Development of Memory-like T Cells during Chronic Infection. Immunity 47, 1129–1141 e1125. 10.1016/j.immuni.2017.11.021.

32. Im, S.J., Obeng, R.C., Nasti, T.H., McManus, D., Kamphorst, A.O., Gunisetty, S., Prokhnevska, N., Carlisle, J.W., Yu, K., Sica, G.L., et al. (2023). Characteristics and anatomic location of PD-1(+)TCF1(+) stem-like CD8 T cells in chronic viral infection and cancer. Proc Natl Acad Sci U S A 120, e2221985120. 10.1073/pnas.2221985120.

33. Leong, Y.A., Chen, Y., Ong, H.S., Wu, D., Man, K., Deleage, C., Minnich, M., Meckiff, B.J., Wei, Y., Hou, Z., et al. (2016). CXCR5(+) follicular cytotoxic T cells control viral infection in B cell follicles. Nat Immunol 17, 1187–1196. 10.1038/ni.3543.

34. Mueller, S.N., Matloubian, M., Clemens, D.M., Sharpe, A.H., Freeman, G.J., Gangappa, S., Larsen, C.P., and Ahmed, R. (2007). Viral targeting of fibroblastic reticular cells contributes to immunosuppression and persistence during chronic infection. Proc Natl Acad Sci U S A 104, 15430–15435. 10.1073/pnas.0702579104.

35. Reinhardt, R.L., Liang, H.E., and Locksley, R.M. (2009). Cytokine-secreting follicular T cells shape the antibody repertoire. Nat Immunol 10, 385–393. 10.1038/ni.1715.

36. Abiko, K., Matsumura, N., Hamanishi, J., Horikawa, N., Murakami, R., Yamaguchi, K., Yoshioka, Y., Baba, T., Konishi, I., and Mandai, M. (2015). IFN-gamma from lymphocytes induces PD-L1 expression and promotes progression of ovarian cancer. Br J Cancer 112, 1501–1509. 10.1038/bjc.2015.101.

37. Garcia-Diaz, A., Shin, D.S., Moreno, B.H., Saco, J., Escuin-Ordinas, H., Rodriguez, G.A., Zaretsky, J.M., Sun, L., Hugo, W., Wang, X., et al. (2017). Interferon Receptor Signaling Pathways Regulating PD-L1 and PD-L2 Expression. Cell Rep 19, 1189–1201. 10.1016/j.celrep.2017.04.031.

38. Bazhin, A.V., von Ahn, K., Fritz, J., Werner, J., and Karakhanova, S. (2018). Interferon-alpha Up-Regulates the Expression of PD-L1 Molecules on Immune Cells Through STAT3 and p38 Signaling. Front Immunol 9, 2129. 10.3389/fimmu.2018.02129.

39. Dahling, S., Mansilla, A.M., Knopper, K., Grafen, A., Utzschneider, D.T., Ugur, M., Whitney, P.G., Bachem, A., Arampatzi, P., Imdahl, F., et al. (2022). Type 1 conventional dendritic cells maintain and guide the differentiation of precursors of exhausted T cells in distinct cellular niches. Immunity 55, 656–670 e658. 10.1016/j.immuni.2022.03.006.

40. Magen, A., Hamon, P., Fiaschi, N., Soong, B.Y., Park, M.D., Mattiuz, R., Humblin, E., Troncoso, L., D’Souza, D., Dawson, T., et al. (2023). Intratumoral dendritic cell-CD4(+) T helper cell niches enable CD8(+) T cell differentiation following PD-1 blockade in hepatocellular carcinoma. Nat Med 29, 1389–1399. 10.1038/s41591-023-02345-0.

41. Boukhaled, G.M., Harding, S., and Brooks, D.G. (2021). Opposing Roles of Type I Interferons in Cancer Immunity. Annu Rev Pathol 16, 167–198. 10.1146/annurev-pathol-031920-093932.

42. Le Bon, A., Durand, V., Kamphuis, E., Thompson, C., Bulfone-Paus, S., Rossmann, C., Kalinke, U., and Tough, D.F. (2006). Direct stimulation of T cells by type I IFN enhances the CD8+ T cell response during cross-priming. J Immunol 176, 4682–4689. 10.4049/jimmunol.176.8.4682.

43. Curtsinger, J.M., Agarwal, P., Lins, D.C., and Mescher, M.F. (2012). Autocrine IFN-gamma promotes naive CD8 T cell differentiation and synergizes with IFN-alpha to stimulate strong function. J Immunol 189, 659–668. 10.4049/jimmunol.1102727.

44. Shaabani, N., Duhan, V., Khairnar, V., Gassa, A., Ferrer-Tur, R., Haussinger, D., Recher, M., Zelinskyy, G., Liu, J., Dittmer, U., et al. (2016). CD169(+) macrophages regulate PD-L1 expression via type I interferon and thereby prevent severe immunopathology after LCMV infection. Cell Death Dis 7, e2446. 10.1038/cddis.2016.350.

45. Wherry, E.J., Blattman, J.N., Murali-Krishna, K., van der Most, R., and Ahmed, R. (2003). Viral persistence alters CD8 T-cell immunodominance and tissue distribution and results in distinct stages of functional impairment. J Virol 77, 4911–4927. 10.1128/jvi.77.8.4911-4927.2003.

46. Shah, K., Al-Haidari, A., Sun, J., and Kazi, J.U. (2021). T cell receptor (TCR) signaling in health and disease. Signal Transduct Target Ther 6, 412. 10.1038/s41392-021-00823-w.

47. Liu, T., Zhang, L., Joo, D., and Sun, S.C. (2017). NF-kappaB signaling in inflammation. Signal Transduct Target Ther 2, 17023-. 10.1038/sigtrans.2017.23.

48. Zhang, W., and Liu, H.T. (2002). MAPK signal pathways in the regulation of cell proliferation in mammalian cells. Cell Res 12, 9–18. 10.1038/sj.cr.7290105.

49. Martinez, G.J., Pereira, R.M., Aijo, T., Kim, E.Y., Marangoni, F., Pipkin, M.E., Togher, S., Heissmeyer, V., Zhang, Y.C., Crotty, S., et al. (2015). The transcription factor NFAT promotes exhaustion of activated CD8(+) T cells. Immunity 42, 265–278. 10.1016/j.immuni.2015.01.006.

50. Seo, H., Gonzalez-Avalos, E., Zhang, W., Ramchandani, P., Yang, C., Lio, C.J., Rao, A., and Hogan, P.G. (2021). BATF and IRF4 cooperate to counter exhaustion in tumor-infiltrating CAR T cells. Nat Immunol 22, 983–995. 10.1038/s41590-021-00964-8.

51. Zak, J., Pratumchai, I., Marro, B.S., Marquardt, K.L., Zavareh, R.B., Lairson, L.L., Oldstone, M.B.A., Varner, J.A., Hegerova, L., Cao, Q., et al. (2024). JAK inhibition enhances checkpoint blockade immunotherapy in patients with Hodgkin lymphoma. Science 384, eade8520. 10.1126/science.ade8520.

52. Mathew, D., Marmarelis, M.E., Foley, C., Bauml, J.M., Ye, D., Ghinnagow, R., Ngiow, S.F., Klapholz, M., Jun, S., Zhang, Z., et al. (2024). Combined JAK inhibition and PD-1 immunotherapy for non-small cell lung cancer patients. Science 384, eadf1329. 10.1126/science.adf1329.

53. Brooks, D.G., McGavern, D.B., and Oldstone, M.B. (2006). Reprogramming of antiviral T cells prevents inactivation and restores T cell activity during persistent viral infection. J Clin Invest 116, 1675–1685. 10.1172/JCI26856.

54. Brooks, D.G., Teyton, L., Oldstone, M.B., and McGavern, D.B. (2005). Intrinsic functional dysregulation of CD4 T cells occurs rapidly following persistent viral infection. J Virol 79, 10514–10527. 10.1128/JVI.79.16.10514-10527.2005.

55. Levine, J.H., Simonds, E.F., Bendall, S.C., Davis, K.L., Amir el, A.D., Tadmor, M.D., Litvin, O., Fienberg, H.G., Jager, A., Zunder, E.R., et al. (2015). Data-Driven Phenotypic Dissection of AML Reveals Progenitor-like Cells that Correlate with Prognosis. Cell 162, 184–197. 10.1016/j.cell.2015.05.047.

56. Weber, L.M., Nowicka, M., Soneson, C., and Robinson, M.D. (2019). diffcyt: Differential discovery in high-dimensional cytometry via high-resolution clustering. Communications Biology 2, 183. 10.1038/s42003-019-0415-5.

57. Bais, A.S., and Kostka, D. (2020). scds: computational annotation of doublets in single-cell RNA sequencing data. Bioinformatics 36, 1150–1158. 10.1093/bioinformatics/btz698.

58. Korsunsky, I., Millard, N., Fan, J., Slowikowski, K., Zhang, F., Wei, K., Baglaenko, Y., Brenner, M., Loh, P.R., and Raychaudhuri, S. (2019). Fast, sensitive and accurate integration of single-cell data with Harmony. Nat Methods 16, 1289–1296. 10.1038/s41592-019-0619-0.

59. Liberzon, A., Subramanian, A., Pinchback, R., Thorvaldsdottir, H., Tamayo, P., and Mesirov, J.P. (2011). Molecular signatures database (MSigDB) 3.0. Bioinformatics 27, 1739–1740. 10.1093/bioinformatics/btr260.

